# Hypothalamic deiodinase type-3 establishes the period of circannual interval timing in mammals

**DOI:** 10.1101/2025.04.22.650143

**Authors:** Calum Stewart, T. Adam Liddle, Elisabetta Tolla, Jo Edward Lewis, Christopher Marshall, Neil P Evans, Peter J Morgan, Fran JP Ebling, Tyler J Stevenson

## Abstract

Animals respond to environmental cues to time phenological events, but the intrinsic mechanism of circannual timing remains elusive. We used transcriptomic sequencing and frequent sampling of multiple hypothalamic nuclei in Djungarian hamster to examine the neural and molecular architecture of circannual interval timing. Our study identified three distinct phases of transcript changes, with deiodinase type-3 (*Dio3*) expression activated during the early induction phase. Subsequent work demonstrated that targeted mutation of *Dio3* using CRISPR-Cas resulted in a shorter period for circannual interval timing. Hamsters that are non-responsive to short photoperiod and fail to show any winter adaptations do not display changes in *Dio3* expression do not show any change in body mass or pelage. Our work demonstrates that changes in *Dio3* induction is essential for setting the period of circannual interval timing.

## Introduction

Phenology of key life history traits is common across plant (Piao et al., 2019) and animal kingdoms (Cohen et al., 2018). The annual changes in daylength are the predominant environmental cue which animals use to time seasonal life history transitions (Liddle et al., 2022; Pérez et al., 2019; Wood and Loudon, 2014). Plants and animals also exhibit endogenous circannual timing in the absence of any change in environmental cues. For example, bird migration (Gwinner and Dittami, 1990), mammalian hibernation (Pengelley and Fisher, 1957), and reproduction (Woodfill et al., 1994; Lincoln et al., 2006) are all driven by robust intrinsically generated *circannual clocks*, the cycle of which nearly match a 12-month period. Some *circannual timers* estimate an interval period (e.g., 6-month) in which programmed changes in physiology and morphology occur in anticipation of the next season. Such interval timers are commonly observed as flowering in plants (Duncan et al., 2015), diapause in insects (Denlinger, 1974) and spring emergence in rodents (Prendergast et al., 2004). Interval timers are typically characterized by having light-dependent induction, maintenance and recovery phases.

Annual mammalian life history transitions are typically associated with major changes in energy demand (Ricklefs, 1991). While the melanocortin system, including neuropeptide y (*Npy*), agouti-related peptide (*Agrp*), and melanocortin receptors are known to regulate short-term changes in energy homeostasis (Yeo et al., 2021), the mechanisms and anatomical structures implicated in long-term, circannual variation in energy rheostasis are not well characterized. Somatostatin (*Sst*) (Marshall et al., 2024; Petri et al., 2016, 2014), proopiomelanocortin (POMC) (Bao et al., 2019; Mercer et al., 2000), and VGF nerve growth factor (Barrett et al., 2005) are known to be strong correlates of seasonal variation in energetic state. Tanycyte somas are essential for the integration of environmental and physiological signals required for circannual interval timing. Tanycytes are localized in the ependymal layer of the third ventricle and are highly sensitive to nutrient state (Bolborea et al., 2020) and receive photoperiodic signaling derived from thyrotropes in the pars tuberalis (Hanon et al., 2008; Wood et al., 2020). Previous work has established deiodinase type-2 (*Dio2*) and type-3 (*Dio3*) expression is anatomically localized to tanycytes and coordinate triiodothyronine-dependent annual transitions in physiological state (Bao et al., 2019; Ebling and Lewis, 2018; Hanon et al., 2008; Murphy et al., 2012; Petri et al., 2016; Wood et al., 2020). Here we delineate the molecular architecture of circannual interval timing by the hypothalamus and pituitary gland and tested the conjecture that *Dio3*, in the Djungarian hamster (*Phodopus sungorus*), is upregulated during the induction of circannual interval timing for energy rheostasis, and functions to establish the duration (or period) of the circannual interval timer.

## Results

Male Djungarian hamsters were either kept in a long photoperiod (LP) control condition or moved from LP to short photoperiod (SP) conditions (Fig.1). Djungarian hamsters remain in LP phenotype unless exposed to SP. Pelage color, torpor and body mass were used to monitor the induction, maintenance, and recovery phases of the circannual timer (Fig.1A and C *SI Appendix* Movies S1 and 2). A full white pelage color was observed between 12-16 weeks after exposure to SP and gradually reversed to the LP agouti color after 28 weeks in SP. Hamsters engaged in torpor between 12-20 weeks in SP (movie S2). Massive, programmed changes in energy rheostasis were observed with a 30% reduction in daily food intake and a 20% decrease in average body mass by 12 weeks SP (Fig.1C and D). Both food intake and body mass started to reverse to LP conditions after 20 weeks in SP. Epididymal adipose tissue mass and plasma insulin concentrations paralleled the change in body mass (Fig. 1D and *SI Appendix* Fig. S1). These reversals in physiological state are indicative of the recovery phase of the circannual interval timer. GLP-1 did not display any change in circulating concentrations (Fig. 1E). Plasma glucose concentration jumped sharply after 16 weeks in SP (Fig. 1E), concurrent with the development of torpor in these animals (movie S2). The observed change in body mass, food intake, adipose tissue and plasma insulin versus the lack of change in GLP-1 highlights the physiological distinction between a programmed rheostatic mechanism characteristic of the circannual interval timing of energy stability in the former, while the latter are driven by short-term homeostatic mechanisms (Stevenson, 2024).

**Figure 1.**
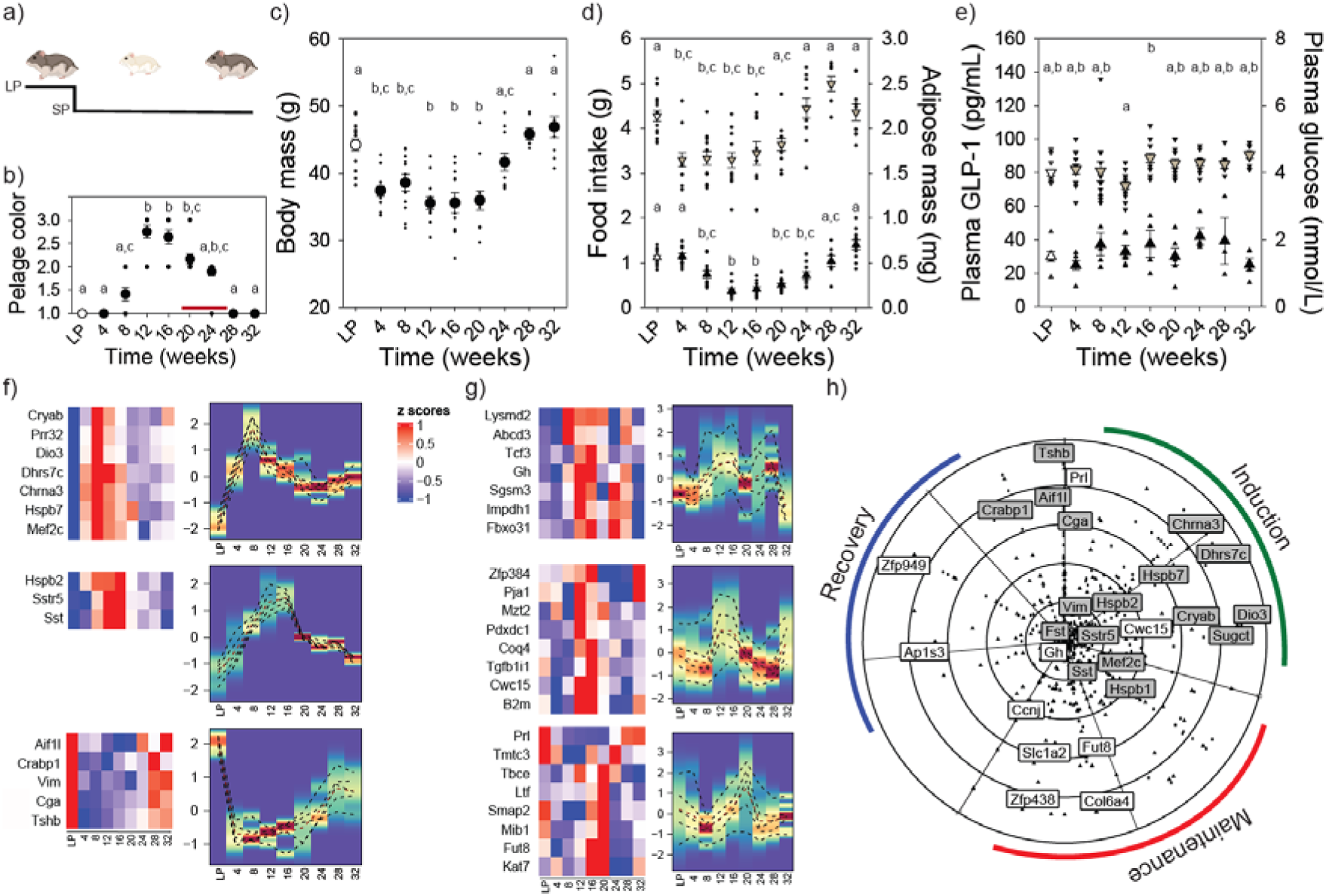
Molecular basis of circannual interval timing for morphology and physiology. Experimental design in which Djungarian hamsters were kept in long photoperiod (LP) or transferred to short photoperiod (SP) (a). SP induced pelage color change from LP agouti to winter white after 12-weeks which reversed to agouti by 28-weeks (F_8,91_ = 50.77; P < 0.001) (b). Torpor was identified in hamster between 12- and 20-weeks SP exposure indicate by red line (b). SP exposure induced significant reduction in body mass (F_8,91_ = 13.428; P < 0.001) (c), food intake (F_8,91_ = 10.860; P < 0.001; denoted as downward triangles) (d) and adipose mass (F_8,91_ = 27.929; P < 0.001; denoted as upward triangles) (d). Plasma GLP-1 did not vary in response to SP manipulation; denoted as upward triangles (e). SP exposure resulted in significant changes in plasma glucose around the onset of torpor; denoted as downward triangles (F_8,91_ = 3.117; P < 0.05) (e). BioDare2.0 heatmaps of mediobasal hypothalamus (f) and pituitary gland (g) transcripts from Djungarian hamster collected at 4-week SP intervals. Transcripts identified as highly rhythmic (FDR < 0.1) showed three distinct phases of expression that coincide with the induction, maintenance and recovery of circannual interval timing. Deiodinase type-3 (*Dio3*) was upregulated during the induction phase, whereas transcripts associated with energy stability (e.g., somatostatin [Sst and Sstr5]) were upregulated during the maintenance phase. All rhythmic transcripts reverted to the LP condition by 28-weeks SP exposure. Polar scatter chart of significant transcripts from mediobasal hypothalamus and pituitary gland provide a comprehensive seasonal clock for mammalian circannual interval timing across neuroendocrine tissues (h). Green line indicates the induction phase, red line indicates the maintenance phase, and the blue line represents the recovery phase. Letters denote significant differences between treatment groups (P < 0.05) (b-e). In panel f and g, the scale bar represents transcript expression as z-scores from 1 (upregulation in red) to -1 (down regulation in blue). Density heatmaps are adjacent to transcript expression heat maps and display the average z-score expression of each individual cluster on the y-axis, and the graph shows percentile lines and density of z-score expression.

We then used Oxford Nanopore transcriptomic sequencing to characterize the molecular changes associated with phases of the circannual interval timer within multiple individual hypothalamic nuclei and the pituitary gland (Fig.1F to G; fig. *SI Appendix* S2 to 6 and Dataset S1 to 4). The development of refractoriness to melatonin, indicative of the recovery phase of the circannual timer, has been shown to develop independently in different hypothalamic regions (Freeman and Zucker, 2001). Molecular data was collected from the mediobasal hypothalamus (Bolborea et al., 2020), dorsomedial hypothalamus (Ebling et al., 2008), and the paraventricular nucleus (Bittman et al., 1991), along with the pituitary gland (Majumdar et al., 2023). This experimental approach provides a high-throughput and high-frequency sampling resolution to comprehensively chart the induction, maintenance and recovery of the mammalian circannual interval timer (Fig. 1H). Biodare2.0 identified 290 (Dataset S1) transcripts in the mediobasal hypothalamus as endogenously rhythmic. Density heat-maps show an early, and robust upregulation of transcripts by 8 weeks SP associated with the circannual interval induction (Fig. 1F). Consistent with previous reports (Petri et al., 2016; Bao et al., 2019; Yoshimura et al., 2003), *Dio3* expression was significantly upregulated and clustered with the initial wave of transcript expression (Fig.1F and *SI Appendix* Fig. S1). A second cluster of transcripts were upregulated during the maintenance phase occurring between 12- and 20-weeks in SP which primarily consisted of somatostatin (*Sst*) and the cognate receptor subtype 5 (*Sstr5*). Increased *Sst* expression coincided with the reduction in body and adipose tissue mass reflecting the conserved role in inhibiting growth and metabolism (*SI Appendix* Figs. S1 and S2). Finally, a third wave of transcripts became upregulated during the recovery phase, and largely resembled the molecular landscape in LP hamsters (Fig.1F). qPCR assays for *Dio3* and *Sst* expression reflected the sequencing transcript count pattern providing independent replication of the sequencing analyses (Fig. S1). In the pituitary gland, 250 transcripts were identified as rhythmic and included genes associated with secretagogue cells such as somatotrophs (*Gh*) and lactotrophs (*Prl*) ( Fig.1g and *SI Appendix* Fig S3 and Dataset S2). Similar rhythmic expression was identified in other hypothalamic nuclei including the paraventricular nucleus (518 transcripts) (*SI Appendix* Fig. S4 and S5 and Dataset S3), and the dorsomedial hypothalamus (374 transcripts) (Fig. S4 and *SI Appendix* S6 and Dataset S4).

### *Dio3*-dependent changes in triiodothyronine induced body mass loss via *Sst* expression

Treatment of Djungarian hamster with the somatostatin agonist pasireotide decreases body mass and is sufficient to drive changes similar to SP exposure (Dumbell et al., 2017, 2015). Further, certain SST receptor subtypes appear to be involved in driving seasonal torpor in Djungarian hamsters (Scherbarth et al., 2015). To determine if hypothalamic *Sst* expression reflects the programmed rheostatic change in energy state governed by circannual interval timing or homeostatic cues, tissues were collected from *ad libitum* (AL) fed hamsters under either LP control condition or the maintenance phase (12-week SP). To induce a negative energetic state, hamsters experienced an acute overnight food restriction (FR; 16hrs) (Fig. 2A). As anticipated, body mass decreased approximately 30% after exposure to SP (Fig.2B, closed symbols); homeostatic negative energy state challenges by food restriction reduced body mass by 1.5g on average for both LP (open symbols) and SP conditions (Fig.2C). Epididymal adipose tissue mass was higher in LP compared to SP but did not decrease in response to overnight FR (*SI Appendix* Fig. S7). Plasma insulin was significantly reduced in response to SP treatment and FR suggesting circulating levels are an output of both rheostatic and homeostatic signals (Fig. S7). The two manipulations permit the ability to dissect the impact of rheostatic changes associated with circannual interval timing from those involved in short-term homeostatic changes in energy balance. Mediobasal hypothalamic *Sst* expression was significantly increased in SP conditions, but insensitive to homeostatic challenges in energy state (Fig.2D). In contrast, mediobasal hypothalamic *Npy* expression was significantly upregulated in food restricted hamsters but was similar across LP and SP conditions (Fig.2E). *Prl* expression was higher in LP compared to SP conditions (Fig. 2F). *Gh* expression did not change in response to either SP induced circannual changes in energy state, or homeostatic manipulations (Fig. S7). To establish whether mediobasal hypothalamic *Sst* expression is regulated by upstream *Dio3*-dependent changes in local thyroid hormone catabolism, SP housed hamsters received sub-cutaneous injections with either vehicle or triiodothyronine. A single triiodothyronine injection in SP hamsters was sufficient to reduce *Sst* expression compared to vehicle controls (Fig. 2G). These data indicate SP induced *Dio3* expression removes T3-dependent inhibition of *Sst* expression leading to long-term inhibition of body mass during the maintenance phase of circannual interval timing of rheostatic energy balance.

**Figure 2.**
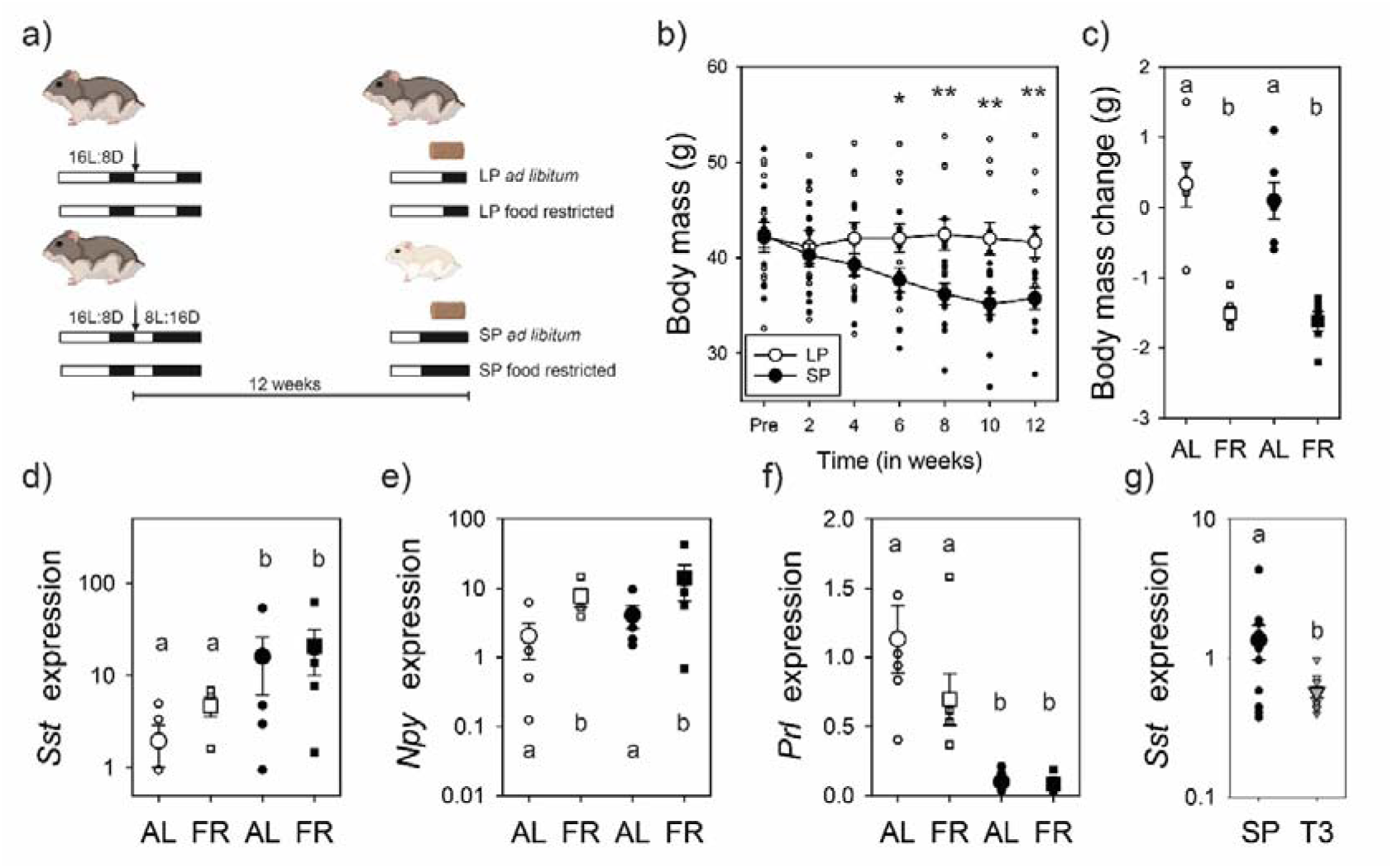
Somatostatin expression reflects programmed circannual interval timing that is dependent on *Dio3* regulation of local triiodothyronine signaling. The experimental design in which SP exposure induced rheostatic reduction in body mass after 12-weeks, and then hamsters experienced either an overnight food restriction (FR) for 16 hours or maintained food ad libitum (AL) (a). SP induced a significant reduction in body mass (F_1,22_ = 3.85; P < 0.01) (b). Food restriction further reduced body mass (F_1,22_ = 25.46; P < 0.001) (c). *Sst* expression in the mediobasal hypothalamus was significantly increased in SP (F_1,15_ = 5.59; P < 0.05) but did not change after manipulations in nutritional availability (d). *Npy* expression was increased after food restriction (F_1,15_ = 6.12; P < 0.05) but did not change with SP exposure (e). Prolactin (*Prl*) expression in the pituitary gland was downregulated in response to SP (F_1,20_ = 28.53; P < 0.001), and insensitive to food restriction. 12-weeks of SP were found to increase *Sst* expression in the hamster mediobasal hypothalamus and levels were significantly reduced in response to a single triiodothyronine (T3) injection (g). *P < 0.05 and **P < 0.01 denote significant difference between SP and LP conditions (b). Letters denote significant difference between treatment groups (P < 0.05) (c-g).

### *Dio3* disfunction reduces the period of circannual interval timing

Next, we sought to establish the functional role of *Dio3* signaling for circannual interval timing of energy rheostasis in hamsters. Targeted genomic mutations of the *Dio3* gene localized to the MBH were achieved via intracerebroventricular (ICV) injection of CRISPR-Cas9 (*Dio3^cc^*; Fig. S8) to assess the functional role for circannual interval timing. Control hamsters were administered a blank crispr-cas9 construct (*Dio3^wt^*). Two weeks following surgery, hamsters were transferred to SP and programmed circannual changes in body mass (Fig.3B) and pelage were monitored (*SI Appendix* Fig. S8). *Dio3^cc^* hamsters had slower body mass loss and regained body mass significantly quicker during the recovery phase, compared to *Dio3^wt^* controls. Similarly, the change in pelage color occurred later in *Dio3^cc^* compared *Dio3^wt^*, and *Dio3^cc^*was never observed to develop full white pelage (*SI Appendix* Fig. S8). Lomb-Scargle period analysis of body mass identified that *Dio3^cc^* hamsters have notably shorter duration (period = 26.959) compared to *Dio3^wt^* (period = 32) (Figs. 3C and D). There was no significant difference in terminal white adipose tissue mass, or daily food intake. A subpopulation of hamsters was physiologically nonresponsive (NR) to the SP manipulation and maintained the higher LP body mass (Fig.3E). *Dio3* expression in the mediobasal hypothalamus of these NR hamsters remained exceptionally low compared to responsive SP hamsters at 8- and 12-weeks following exposure to SP (Fig.3F). These data suggest that the inability to increase *Dio3* expression in NR hamsters acts as a form of naturally occurring disfunction that prevents SP induction of circannual interval timing.

**Figure 3.**
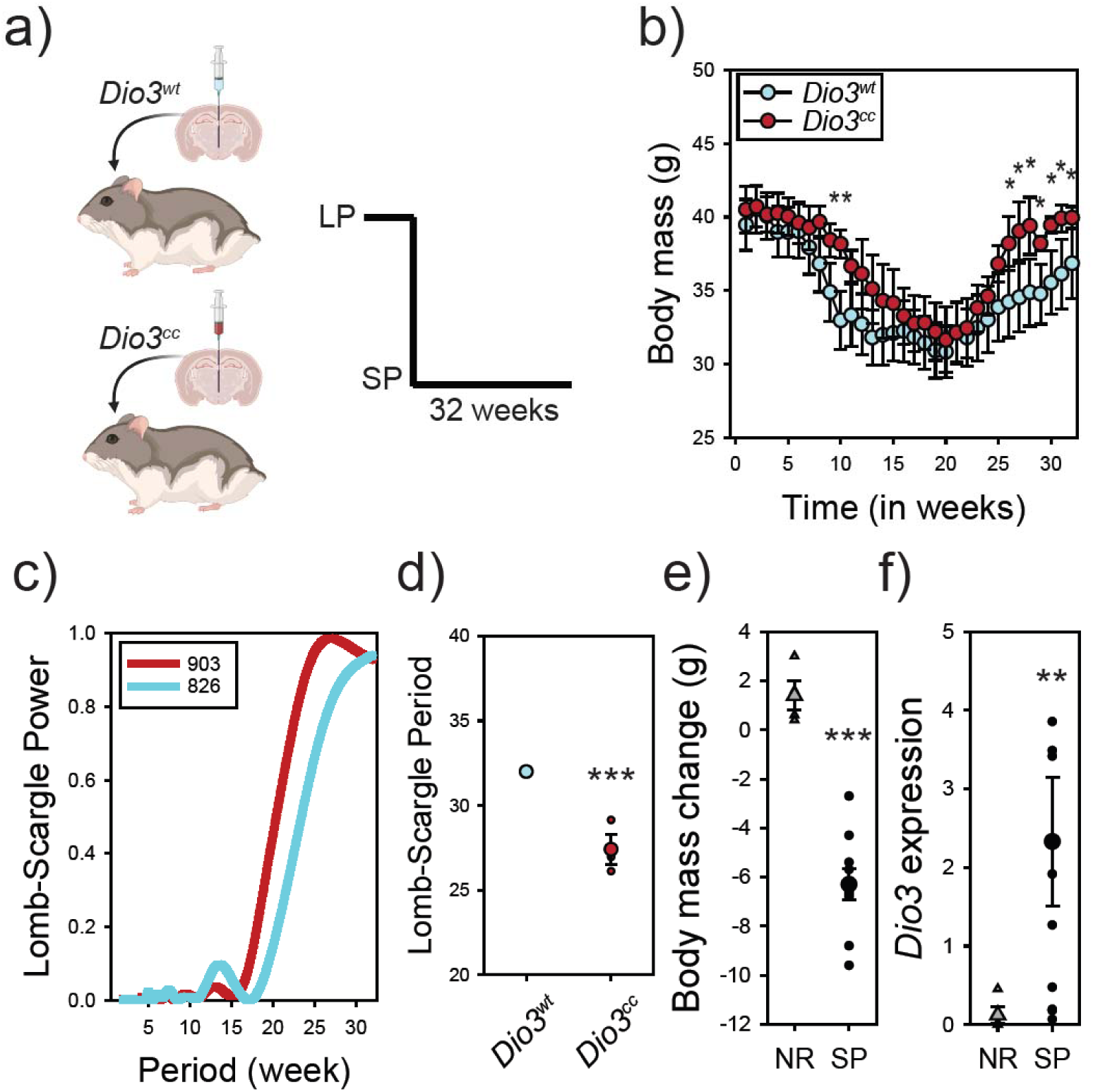
*Dio3* functions to time circannual interval duration in hamsters. Hamsters received intracerebroventricular injections to target *Dio3* expressing cells in the tanycytes localized to the mediobasal hypothalamus (a). Crispr-Cas9 constructs were packaged into lentiviral vectors to generate blank control hamsters (*Dio3^wt^*) or contain gRNAs that mutated the *Dio3* gene (*Dio3^cc^*). Hamsters were then exposed to SP conditions and the circannual interval timer was assessed by monitoring body mass (b). *Dio3^cc^* hamsters were slower to initiate interval timing as evidenced by higher body mass at 10-weeks SP exposure (P < 0.001) and recovered body mass quicker at 26 to 32 (P < 0.05) weeks. Lomb-Scargle analyses identified a significant reduction in the period of the circannual interval timer (Welch’s t-test; t_3_ = 2.62; P < 0.05) (c,d). Example *Dio3^cc^* (hamster #903) and *Dio3^wt^* (hamster #825) are presented in (c). A subpopulation of hamsters did not decrease body mass in response to SP exposure and termed non-responsive (NR) (t_12_ = 7.12; P < 0.001) (e). *Dio3* expression in the mediobasal hypothalamus of NR hamster was nearly nondetectable after exposure to 8-12 weeks of SP exposure compared to SP control hamsters (t_12_ = -3.78; P < 0.005) (f). *P < 0.05, **P < 0.01, and ***P < 0.001.

## Discussion

The molecular and cellular basis of circannual clocks and timers in vertebrates is not well characterized (Helm and Stevenson, 2014). Here, we sequenced the transcriptomes of three hypothalamic regions and the pituitary gland, key neuroendocrine structures which govern programmed circannual rheostatic and homeostatic processes. Our approach provided a high sampling frequency of the circannual interval timer in Djungarian hamsters. The findings produced a robust and comprehensive neural and molecular database which facilitates the delineation of a circannual interval timer in mammals. Several novel transcripts displayed a close correlation with the induction phase of the photoperiodic response. Among these was *Dio3*, conducive with the removal of thyroid hormone (e.g., thyroxine, triiodothyronine) being a key step in the transition to the winter phenotype. The photoinduced change in *Dio3* expression induces a long-term change in physiology and morphology and our results demonstrate that it is insensitive to short-term alterations in response to other environmental cues (e.g., nutritional state). Functional manipulation of *Dio3* expression also delayed the induction and enhanced the reversal of the winter phenotype suggesting that upregulation of *Dio3* acts to establish the period of circannual interval timing.

The transcriptomic data are consistent with evidence that shows *Sst* expression in the arcuate nucleus is one critical event for the maintenance of a reduced rheostatic energy state during the winter season (Marshall et al., 2024; Petri et al., 2016, 2014). In hamsters, *Sst* expression is upregulated after *Dio3* induction and is associated with the low body mass from week 12-20 before a downregulation in transcript levels coinciding with increased body mass. By increasing circulating triiodothyronine in hamsters maintained in SP conditions, our manipulation indicates that elevated hormone concentrations lead to the reduction in *Sst* expression in the mediobasal hypothalamus (Fig.2). The overall expression patterns provide the ability to develop a linear series of steps in which LP conditions have high local concentrations of triiodothyronine in the mediobasal hypothalamus (Helfer and Stevenson, 2020; Murphy et al., 2012) which reduces the levels of *Sst* expression resulting in consistently higher body mass. Exposure to SP induces *Dio3*, which catabolizes triiodothyronine leading to the removal of the inhibition of *Sst* expression and reduction in body mass. Local synthesis of triiodothyronine in the mediobasal hypothalamus by tanycytes is an evolutionary conserved process for timing photoperiod-induced transitions across the animal kingdom (Ebling and Lewis, 2018; Lewis and Ebling, 2017). Photoperiodic signals derived from the pars tuberalis via thyrotropin-stimulating hormone are essential to regulate *Dio3* (and *Dio2*) expression in tanycytes leading to increased triiodothyronine content in LP conditions for mammals (22,23), birds (Nakao et al., 2008) and fish (Lorgen et al., 2015). The current limitation is understanding how SP signals regulate *Dio3* expression. Undoubtedly, the duration of melatonin secretion during the daily nocturnal phase is essential (Helfer et al., 2019). Future research is necessary to identify the conserved upstream signals that govern the timing of *Dio3* expression.

These data support the existence of a neuroendocrine pathway for the long-term rheostatic regulation, and another for short-term homeostatic control of energy stability (Stevenson, 2024). The rheostatic pathway consists of triiodothyronine (T3) signaling dependent on a *Dio3* switch to regulate *Sst* expression in the mediobasal hypothalamus whereas short-term regulation of energy stability via nutrient availability is homeostatically regulated by well-established orexigenic (e.g., *Npy*) (Yeo et al., 2021). In response to the output of both pathways, peripheral effects may be mediated by acute and chronic changes in pancreatic insulin secretion (Fig.4). Overall, the findings provide a clear neural and cellular circuit for circannual interval timing of energy rheostasis and demonstrate *Dio3* as a gene critical for the control of seasonal life history transitions.

**Fig 4.**
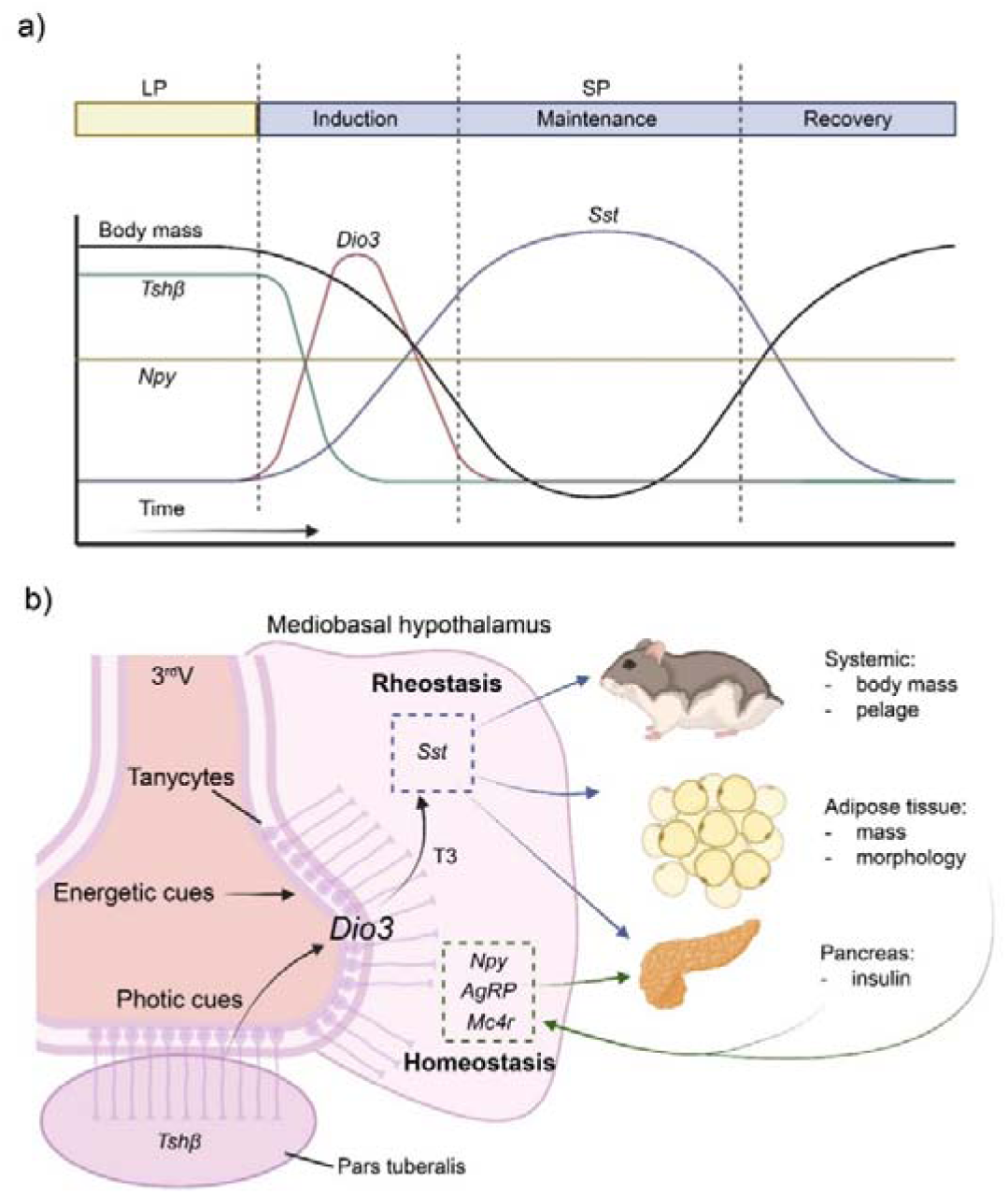
Schematic model for the mechanisms of circannual interval timing. (a) Hamsters held in long photoperiod (LP) have high body mass, thyrotropin-stimulating hormone beta (*Tsh*β) and low somatostatin (*Sst*) and deiodinase type-3 (*Dio3*). Transfer to short photoperiod (SP) results in the induction of circannual interval timing characterized by a reduction in body mass, low *Tsh*β in the pars tuberalis and increased *Dio3* expression in tanycytes. The maintenance phase is associated with increased *Sst* expression which serves to inhibit growth. The onset of the recovery phase is associated with a complete reversal in whole organismal physiology to the LP phenotype. (b) tanycytes along the third ventricle in the mediobasal hypothalamus integrate photic cues derived from *Tsh*β and are sensitive to nutritional cues. The triiodothyronine (T3) output signal from tanycytes regulates *Sst* expression leading to long-term programmed rheostatic changes in body mass that serve to control circannual interval timing in multiple physiological systems. Conversely, homeostatic stability is maintained despite large scale seasonal rhythms in body mass. Homeostatic energy balance is established by well characterized circuits that include neuropeptide Y (*Npy*), agouti-related peptide (*Agrp*) and the melanocortin receptor 4 (*Mc4r*).

## Materials and Methods

### Ethics

All procedures were in accordance with the National Centre for the Replacement, Refinement and Reduction of Animals in Research ARRIVE guidelines (https://www.nc3rs.org.uk/revision-arrive-guidelines). All procedures were approved by the Animal Welfare and Ethics Review Board at the University of Glasgow and conducted under the Home Office Project License PP5701950.

### Animals

All animals were obtained from a colony of Djungarian hamsters (3-8 months of age) kept at the University of Glasgow, Veterinary Research Facility. Animals were held at a room temperature of 21°C, and a humidity of 50%. The colony room was kept on a long photoperiod (16L:8D). All hamsters had food (SDS [BK001]) and water *ad libitum*.

### Experiment 1 – Defining the circannual interval timer in hamsters

Adult male hamsters (N = 100) were required to assess the physiological and molecular basis of circannual interval timing in hamsters. A photoperiod reference group was kept on LP (n=14) (8D:16L) conditions for the duration of the study. The SP treatment groups consisted of hamsters moved to SP (16D:8L) at 4 weeks intervals. The groups consisted of SP for 4 (n=12), 8 (n=12), 12 (n=12), 16 (n=11), 20 (n=12), 24 (n=10), 28 (n=7) and 32 (n=10) weeks. This photoperiod schedule reliably captures the induction, maintenance, and recovery of circannual interval timing in hamsters (Prendergast et al., 2002). Pelage score was determined using visual assessment and defined scale in which a score of 1 is the summer dark agouti coat; a score of 2 is a transition between agouti and white; and a score of 3 is a full winter white pelage (Marshall et al., 2024). Body mass was monitored to the nearest 0.1g using an Ohaus portable scale (TA301). Food intake was measured by weighing the food pellets at the mid-point on two consecutive days to obtain a daily value. Food weight was measured using an Ohaus scale (TA301). Hamsters were killed between 2-4hr after lights on by cervical dislocation followed by exsanguination. Adipose tissue was immediately dissected and measured using a Sartorius cp64 anatomical balance. A terminal blood sample was collected to measure glucose (Accu-check Performa nano blood glucose meter). Circulating levels of insulin and total glucagon-like peptide 1 (GLP1) using ELISA (MesoScale Discovery, UK) at Core Biochemical Assays laboratories, Cambridge, UK. Brain and pituitary gland tissue was quickly extracted and frozen on powdered dry ice and stored at -70°C. Epididymal white adipose tissue was dissected and weighed using Sartorius cp64 anatomical balance and stored at -70°C.

Brains were cut into 200µm coronal sections using a Leica CM1520 cryostat. Anatomical structures (optic tract to the infundibular stem; Bregma -2.12mm to -3.80mm) were used to isolate the mediobasal hypothalamus (MBH), dorsomedial hypothalamus (DMH) and the paraventricular nucleus (PVN). Bilateral tissue punches were performed using an integra Miltex 1mm disposable biopsy punch. Tissue punches were stored at -70°C until transcriptomic sequencing and confirmatory qPCR analyses (see below).

### Experiment 2 – Dissociating mechanisms that govern circannual rheostatic versus homeostatic energy stability

A total of N=35 adult male hamsters were used to determine molecular signatures of rheostatic versus homeostatic energy stability. A LP control group was kept on long photoperiod for the duration of the study (n = 18). A subset of hamsters was moved to SP (n = 17) for 12 weeks to obtain animals in the maintenance phase of circannual interval timing. Food (SDS [BK001]) and tap water were provided *ad libitum* to both LP and SP animals for the 12 weeks duration. Body mass was measured every two weeks using a Traveler Ohaus portable balance. On the final night of the experiment, 50% of hamsters (i.e., LP = 9, SP = 9) were kept on food and water *ad libitum*. The other 50% of hamsters (i.e., LP = 9, SS = 8) served as the homeostatic treatment group and received an acute food restriction by removing all food. Overnight food restriction induces a robust negative energy state that reliably results in a 1–2-gram loss in hamster body mass (Bao *et al*., 2019). Body mass was measured prior to food restriction, and again before tissue collection on the subsequent day. Hamsters were killed between 2-4hr after lights on by cervical dislocation and then exsanguination. A terminal blood sample was collected to measure insulin (MesoScale Discovery, Core Biochemical Assays laboratories, Cambridge, UK). Body mass and epididymal white adipose tissue mass were measured using a Sartorius cp64 anatomical balance. Brains and pituitary glands were dissected and were immediately placed on dry ice and stored at -70°C. Brains were cut into 200µM sections using a Leica CM1520 cryostat. Anatomical structures (optic tract to the infundibular stem; Bregma -2.12mm to -3.80mm) were used to isolate the MBH. Bilateral tissue punches were performed using an integra Miltex 1mm disposable biopsy punch. Brain tissue punches and pituitary gland samples were stored at -70°C until qPCR assay (see below).

### Experiment 3 – Sufficiency of triiodothyronine to induce LP neuroendocrine state

Adult male hamsters (n = 35) were used to determine the impact of triiodothyronine on *Sst* expression. Hamsters were transferred from LP to SP for 12 weeks (n = 24) to initiate circannual interval timing and establish the maintenance phase. Hamsters were divided into two groups for the last week of SP treatment. A SP reference group received subcutaneous injections of 0.9% w/v saline for 7 days (n = 10). To determine the sufficiency of triiodothyronine to regulate *Sst* expression, hamsters (n=11) were administered a single subcutaneous injection of 5µg/100ul triiodothyronine (Merck 102467157) prior to lights off on the final night. Subcutaneous injections of triiodothyronine are well established to regulate transcript expression in the hamster hypothalamus (Banks et al., 2016; Bao et al., 2019; Stevenson et al., 2014). The subsequent day hamsters were sacrificed by cervical dislocation followed by exsanguination between 4-5hr after lights on. Terminal body mass and epididymal adipose tissue mass were recorded using a Sartorius cp64 anatomical balance. Brains were dissected and stored immediately at -70°C until sectioning. Brains were cut into 200µM sections using a Leica CM1520 cryostat. Anatomical structures (optic tract to the infundibular stem; Bregma -2.12mm to -3.80mm) were used to isolate the MBH. Bilateral tissue punches were performed using an integra Miltex 1mm disposable biopsy punch. Tissue punches were stored at -70°C until qPCR assays (see below).

### Experiment 4 – Functional manipulation of *Dio3* expression in hamsters

Targeted neuroanatomical injections, using highly precise Guide RNA (gRNA) against *Dio3* and Crispr-Cas9 vectors was essential to ensure the long-term inhibition of *Dio3* function in the MBH in hamsters; an approach necessary to reliably examine the functional significance of *Dio3* for the duration of circannual interval timing. Custom-made Crispr-cas9 constructs were packaged in a lentiviral vector by Merck Life science UK. Lentiviral vectors are established to effectively transfect in adult Syrian hamsters (Gao M *et al*., 2014) and in Siberian hamsters (Munley K *et al*., 2022). The lentiviral vector used was U6-gRNA:ef1a-puro-2A-Cas9-2A-tGFP (Sigma All-in-One vector). gRNA for *Dio3* were designed using CHOPCHOP (Labun *et al.,* 2019). Three gRNA were designed to target the *Mus musculus Dio3* gene (gRNA1: CGACAACCGTCTGTGCACCCTGG; gRNA2: GTTCCCGCGCTTCCTAGGCACGG, and gRNA3: GACCCAGCCGTCGGATGGGTGGG). Only gRNA 1 and 2 were identified to align with 5’ end of the Siberian hamster *Dio3* gene and were predicted to align with chromosome 12 at locations 110279543 and 110279375 respectively (Fig. S8). To check specificity of gRNA sequences, we conducted BLAST using the NCBI dataset. Both gRNA1 and gRNA2 were found to align 100% with *Dio3* and did not align 23/23 with any other region.

Adult male hamsters (n = 10) were selected to assess the impact of *Dio3* genome modification on circannual interval timing. Hamsters were kept on LP conditions to maintain the summer phenotype. Hamsters were then divided into two treatment groups. The control group included hamsters (n=6) that received intracerebroventricular (ICV) injections of blank Crispr-Cas9 constructs that were packaged into a lentivirus (*Dio3^wt^*; U6-gRNA:ef1a-puro-2A-Cas9-2A-tGFP (Sigma All-in-One vector). The treatment group consisted of hamsters (n=4) that received an ICV injection of Crispr-cas9 constructs that harbored both gRNA1 and gRNA2 (*Dio3^cc^*).

Intracerebroventricular injections provide the ability to induce neuroanatomically localized manipulations in *Dio3* gene function in the adult brain. Crispr-Cas9 constructs were delivered via stereotaxic ICV injection into the third ventricle to target the MBH tanycytes. Under general anesthesia (5% induction, 2% maintenance of isoflurane mixed with oxygen) a SGE series II syringe was positioned along the midline at Bregma -1.5mm, to a depth of -5.7mm. The anatomical coordinates were refined based on estimates taken from adult hamster brains (Steward et al., 2003) and tested using fresh hamster cadavers. Analgesia was administered subcutaneously (5mg/kg Rimadyal and 0.1mg/kg buprenorphine). A small ∼1cm incision was made by scalpel to expose the skull and a small hole was drilled using a Bone Micro Drill for Brain surgery (Harvard Apparatus). Lentivirus was administered using a Pump 11 Elite Programmable Syringe injection system (Harvard Apparatus) and 1ul of viral vector (2-2.5 E+13 vg/ml) was delivered over 10 minutes at a rate of 0.1ul per minute. To allow viral diffusion and prevent backflow the syringe was kept in place after injection for 1-minute and then the syringe was raised by 1mm and kept in place for another 1-minute.

Animals were then moved to heated home cages and given mashed food to aid recovery. Hamsters were provided with 2-weeks recovery and were monitored to ensure stable body mass, locomotor activity and continued food and water intake.

To assess the impact of *Dio3^cc^* on circannual interval timing hamsters were moved to SP for 32 weeks and body mass and pelage were measured biweekly. Pelage score was determined using visual assessment and defined scale in which a score of 1 is the summer dark agouti coat; a score of 2 is a transition between agouti and white; and a score of 3 is a full winter white pelage (Marshall et al., 2024). Body mass was monitored to the nearest 0.1g using an Ohaus portable scale (TA301). After 32 weeks, hamsters were killed 4-5hr after lights on by cervical dislocation followed by exsanguination. Brains were dissected and stored immediately at -70°C. Brains were cut into 200µM sections using a Leica CM1520 cryostat. Anatomical structures (optic tract to the infundibular stem; Bregma -2.12mm to -3.80mm) were used to isolate the MBH. Bilateral tissue punches were performed using an integra Miltex 1mm disposable biopsy punch. Tissue punches were stored at -70°C. DNA was extracted from dissected tissues using the DNeasy Blood & Tissue Kit (QIAGEN) as per manufacturer’s instructions. The *Dio3* gene was amplified using OneTaq Quick-Load 2X mastermix, as per manufacturer’s instructions (New England Biolabs). Amplification was achieved using a SimpliAmp thermal cycler (Applied Biosystems) the following thermal cycling conditions (94°C for 30 seconds, (94°C for 15 seconds, 62°C for 30 seconds, 68°C for 3 minutes) for 45 cycles, 68°C for 5 minutes, 4°C until further analysis.). The resultant amplicon was purified using the Qiaquick PCR purification kit (Qiagen) as per manufacturer’s instructions. Isolated *Dio3* PCR amplicons were sequenced using the Eurofins LightRun sanger sequencing service. The forward primer provided to Eurofins was CATGCTCCGCTCCCTGCTGCTTCA (5’-3’). CRISPR modified transcripts were aligned to a reference wild type transcript (animal #821) using the Tracy sanger basecaller and aligner (Rausch et al., 2020). Alignments were visualized using the Ugene integrated bioinformatics tool (Okonechnikov et al., 2012). In case mutation was not obvious, an additional analysis was carried out using the associated online resource SAGE (https://www.gear-genomics.com/), to investigate the decomposition error of the modified transcripts with the wild-type control (821). Four *Dio3^wt^* hamsters had *Dio3* sequences that fully matched the reference genome (Fig. S8c). Two hamster *Dio3* sequences did not automatically align, and the raw reads were visually inspected. No mutations or sequence mutations were detected in the raw sequence trace. Three *Dio3^cc^* treated hamsters had *Dio3* sequences with evidence of genomic mutation evidenced by low decomposition error (*SI Appendix* Figs. S8C and D). One *Dio3^cc^*treated hamster did not show any genomic mutation (905) and was removed from the final analysis. Hamster 905 had a body mass circannual interval period of 32 weeks and we propose the animal represents a false positive control.

### Experiment 5 – Naturally occurring variation in *Dio3* expression and photoperiodic response

A subset of hamsters does not physiologically respond to SP treatment and are classified as photoperiodically non-responsive (NR) (Przybylska-Piech and Jefimow, 2022). Adult hamsters (n = 14) were held in LP or SP for 8 weeks. A group of male non responders (n = 4) were identified due to the absence of body mass and pelage change in response to SP (Fig. 3e). A reference group of male SP responders (n = 10) were collected. MBH was extracted, RNA isolated and cDNA synthesized. qPCR was carried out for *Dio3* and *Sst* expression. Hamsters were killed between 2-4hr after lights on.

### RNA extraction

RNA was extracted using the Qiagen RNEasy plus mini kit. Tissues were homogenized using a Polytron PT 1200 E. RNA was then extracted from homogenized tissue following the manufacturer’s instructions. RNA was tested for quality and quantity using an ND-1000 Nanodrop.

### Next Generation Sequencing and data processing

Oxford Nanopore Sequencing was used to carry out transcriptomic sequencing. RNA from MBH, DMH and PVN samples was sequenced using the PCR-cDNA barcoding kit following manufacturer’s instructions (SQK-PCB109; Oxford Nanopore). RNA from the pituitary gland samples was synthesized using the direct cDNA sequencing kit following manufacturer’s instructions (SQK-DCS109 with EXP-NBD104; Oxford Nanopore). Bioinformatic analysis was carried out on Linux within a conda environment. Raw reads (Fast5) were demultiplexed and basecalled using guppy basecaller (4.2.1). Porechop (0.2.4) was used to remove adapters from reads and Filtlong (v0.2.0) was used to filter for quality and read length (length: >25 base pairs, quality score > 9). Transcripts were aligned to Mus musculus genome (GRCm39) using Minimap2 (Li, 2018). Previously, rodent genomes have been demonstrated to show significant similarity, such that cross-species transcriptomic analysis appears feasible (Przybylska-Piech and Jefimow, 2022). Transcript expression levels were generated using Salmon (v0.14.2) (Patro et al., 2017) and EdgeR (v3.24.3) (Robinson et al., 2010) was used to filter lowly expressed transcripts.

### cDNA synthesis and qPCR

RNA was transformed into cDNA for qPCR analysis using superscript III (Invitrogen) as per manufacturer’s instructions. Quantification of transformed cDNA was achieved using Brilliant II SYBR Mastermix (Agilent). Stock forward and reverse primers (100pmol/μl) were mixed and diluted in nuclease free water to 20pmol/μl. A working SYBR mixture was prepared by mixing 1-part primer mixture per 24-part SYBR mastermix. Reaction mixtures were prepared in wells on a 96-well plate by mixing 4.8μl of normalised sample and 4.8μl of SYBR working mix. All reactions were performed in duplicate. Primer and qPCR parameters are outlined in Table S1. qPCR reactions were carried out in a Stratagene Mx3000P thermal cycler. Cycling conditions utilized were in sequence; at 95°C for 5 minutes (denaturing), 40 cycles at (95°C for 30 seconds, X°C for 1 minute – see table, 72°C for 30 seconds with fluorescent measurement at end), 95°C for 1 minute, 55°C for 30 seconds increasing to 95°C (Melt curve analysis). Melt cure analysis was used to determine specificity of amplification. Data was analyzed for meanCT, efficiency and variability using PCR Miner (Zhao and Fernald, 2005). Logfold CT was calculated using the ΔΔCT method using 18s and/or *Hrpt* reference transcripts as these transcripts are stable across photoperiodic conditions (Stewart et al., 2022).

## Statistical Analysis

Raw data is provided in Data file 1 and sequencing data is provided in Table S1-S4. Physiological and qPCR data sets were tested for normality using the Shapiro test and transformed by log transformation if normality was violated. Statistical significance was determined using a two-way ANOVA. Post hoc analysis of significant data was achieved by t-test. Type 1 error was minimized by utilizing the Bonferroni adjustment on post-hoc analyses. Data were analyzed using non-linear regression for rhythmicity using the online resource BioDare 2.0 (Zielinski et al., 2014) (biodare2.ed.ac.uk). The empirical JTK_CYCLE method was used for detection of rhythmicity and the classic BD2 waveform set was used for comparison testing. Rhythmicity was determined by a Benjamini-Hochberg controlled false discovery rate (BH corrected FDR < 0.1). The false discovery rate of this data set was controlled by applying the Benjamini-Hochberg method (Benjamini and Hochberg, 1995). Clusters within the data set were identified using the gap-statistic and clustered using k-means clustering. Gene ontology analysis was carried out using ShinyGO v0.77 (Ge et al., 2020).

## Data and materials availability

All data are available in Dataset S1-S5. The code is available via github repository (https://github.com/CS-Gla/Hamster_seasonal_sci_cs_2024). Sequencing information was deposited in GEO (GSE274003).

## Acknowledgments

The authors thank Gerald Lincoln for helpful discussions which contributed to the conceptualization of the research. The authors thank the technical assistance provided by Nicola Munro, Ana Monterio, and Fallon Cuthill. https://Biorender.com was used for icons used in Fig. 1-3 and Fig. S9. Funding was provided via Leverhulme Trust award LT-RL-2019-06 (TJS) and ISSF Wellcome Trust award (TJS, PJM, FJPE).

## Supplementary Figures

**Fig. S1.**
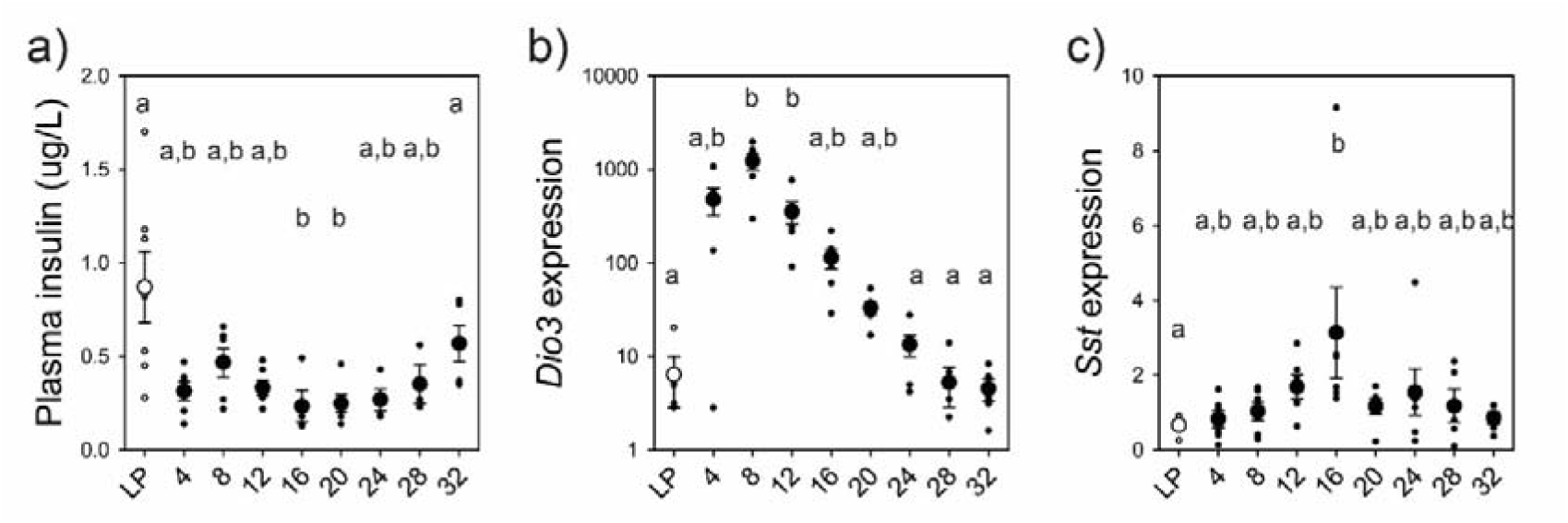
Plasma insulin reflects changes in transcript expression. (A) Plasma insulin of Siberian hamsters across 32-weeks of short photoperiod exposure (H_8_ = 20.401, P = 0.009). (B) Expression of *Dio3* from the MBH of Siberian hamsters, assessed by qPCR, across 32-weeks of short photoperiod exposure (H_8_ = 39.771, P < 0.001). (C) Expression of *Sst* from the MBH of Siberian hamsters, assessed by qPCR, across 32-weeks of short photoperiod exposure (H_8_ =16.002, P = 0.042). Letters denote significance between groups.

**Fig. S2.**
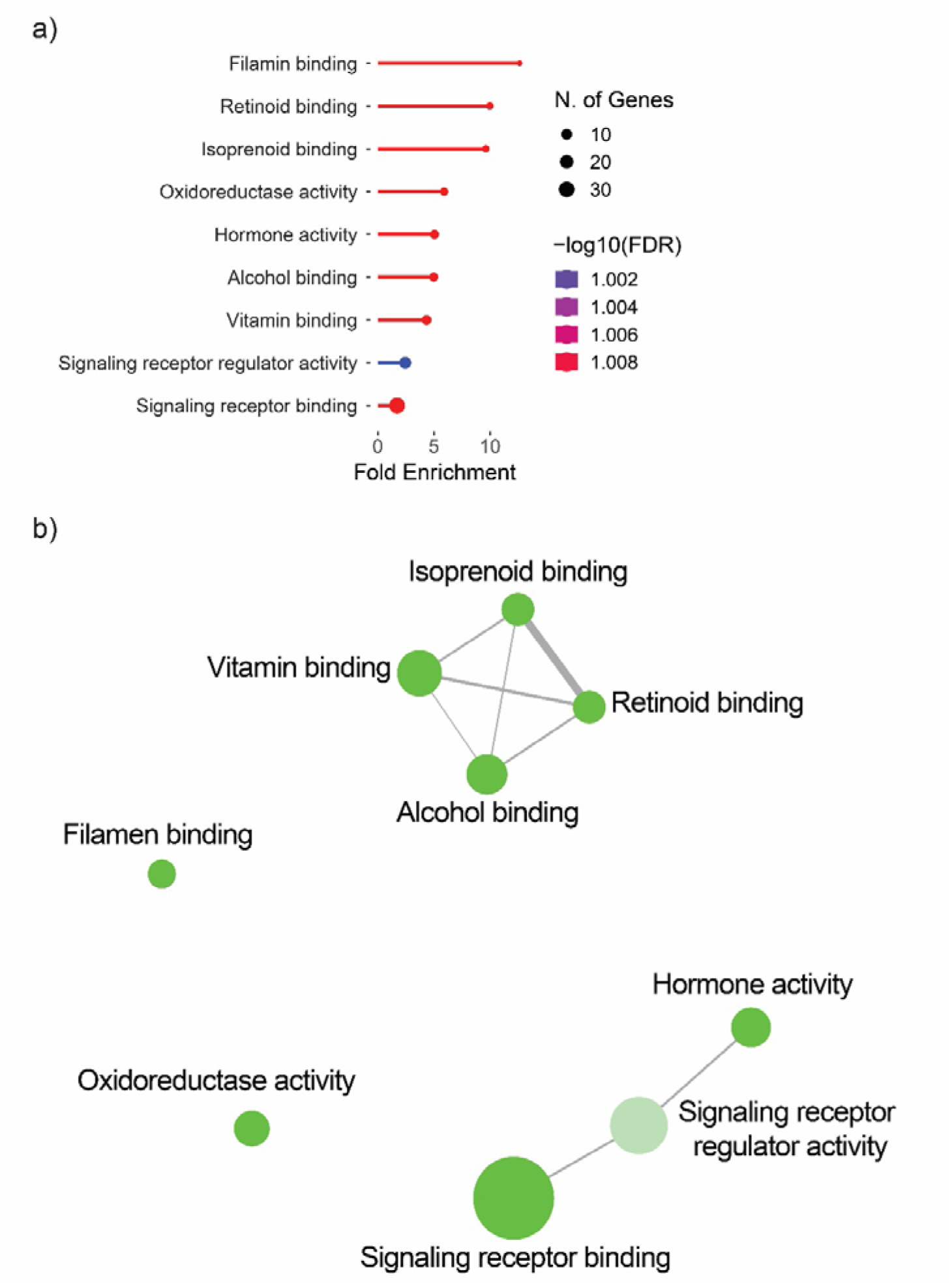
Gene ontology analysis of MBH sequencing reveals well known seasonal pathways. (A) Gene ontology enrichment analysis of significant transcripts from sequencing of Djungarian hamster MBH across 32-weeks of short photoperiod exposure. (B) Interaction network of enriched gene ontology terms. Size of individual points represent the number of transcripts in each term. Thickness of interactions represents shared transcripts between terms.

**Fig. S3.**
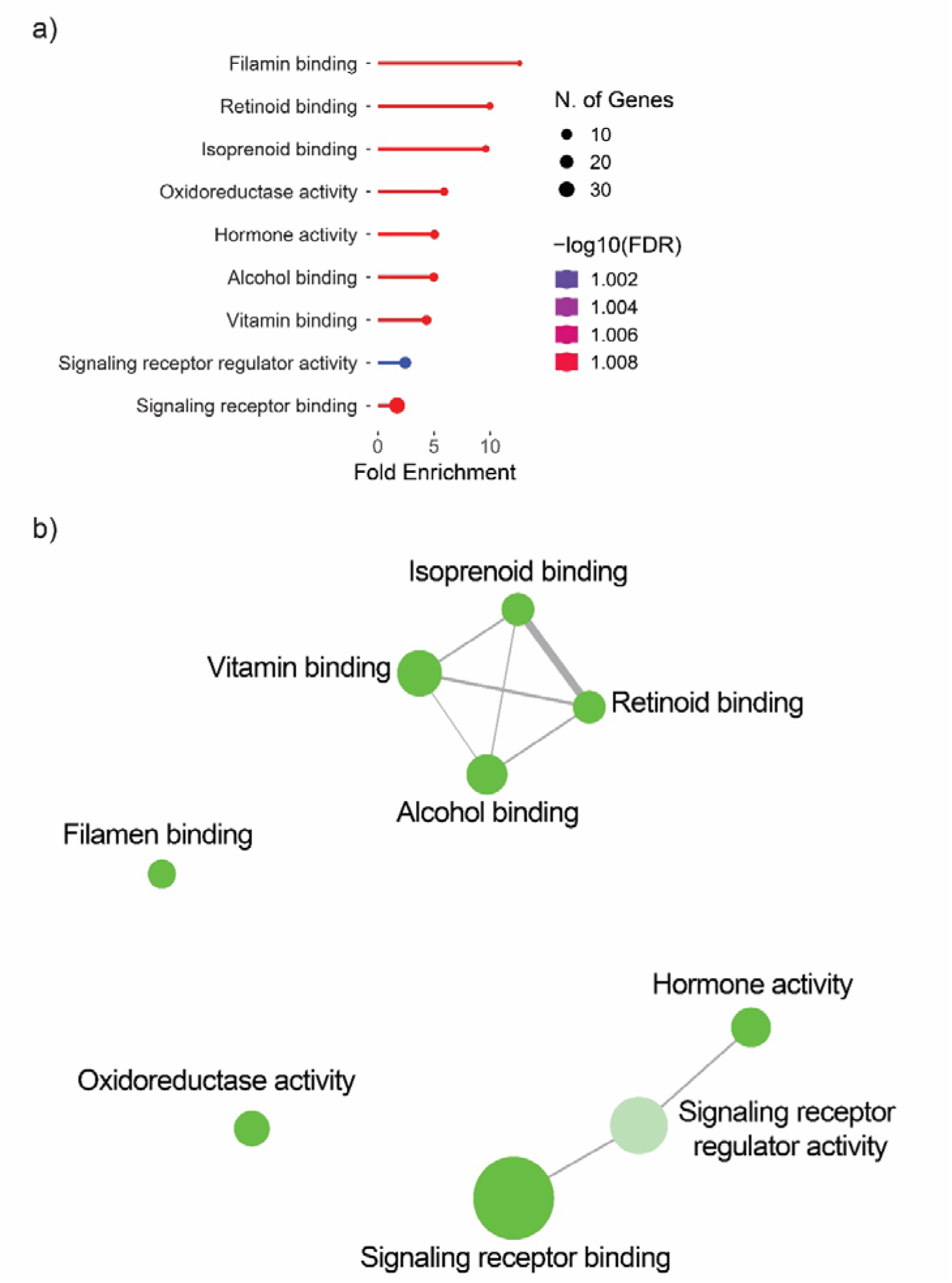
Gene ontology analysis of pituitary gland unveils potential mechanisms for seasonal changes in protein processing and release. (A) Gene ontology enrichment analysis of significant transcripts from sequencing of Djungarian hamster pituitary gland across 32-weeks of short photoperiod exposure. (B) Interaction network of enriched gene ontology terms. Size of individual points represents the number of transcripts in each term. Thickness of interactions represents shared transcripts between terms.

**Fig. S4.**
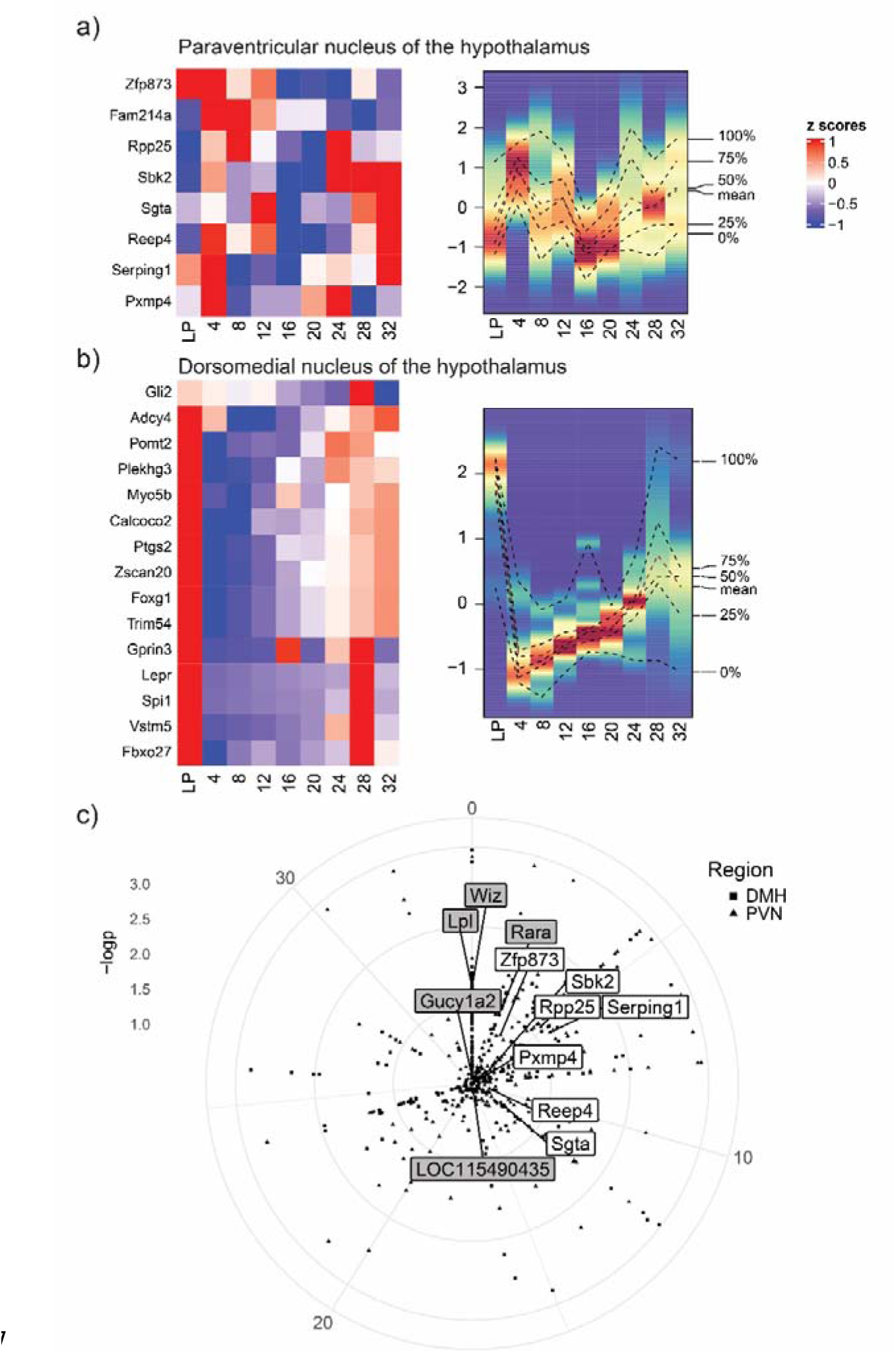
Paraventricular and dorsomedial hypothalamus sequencing unveils novel transcripts and widespread seasonal interval timing within the hypothalamus. (A-B) Heatmaps of significant (FDR < 0.1) Djungarian hamster transcripts displaying sine or cosine rhythmicity were selected from (A) paraventricular hypothalamus and (B) dorsomedial hypothalamus. (C) Polar scatter chart of significant transcripts from Djungarian hamster paraventricular and dorsomedial hypothalamus, displaying peak of expression and -log(FDR).

**Fig. S5.**
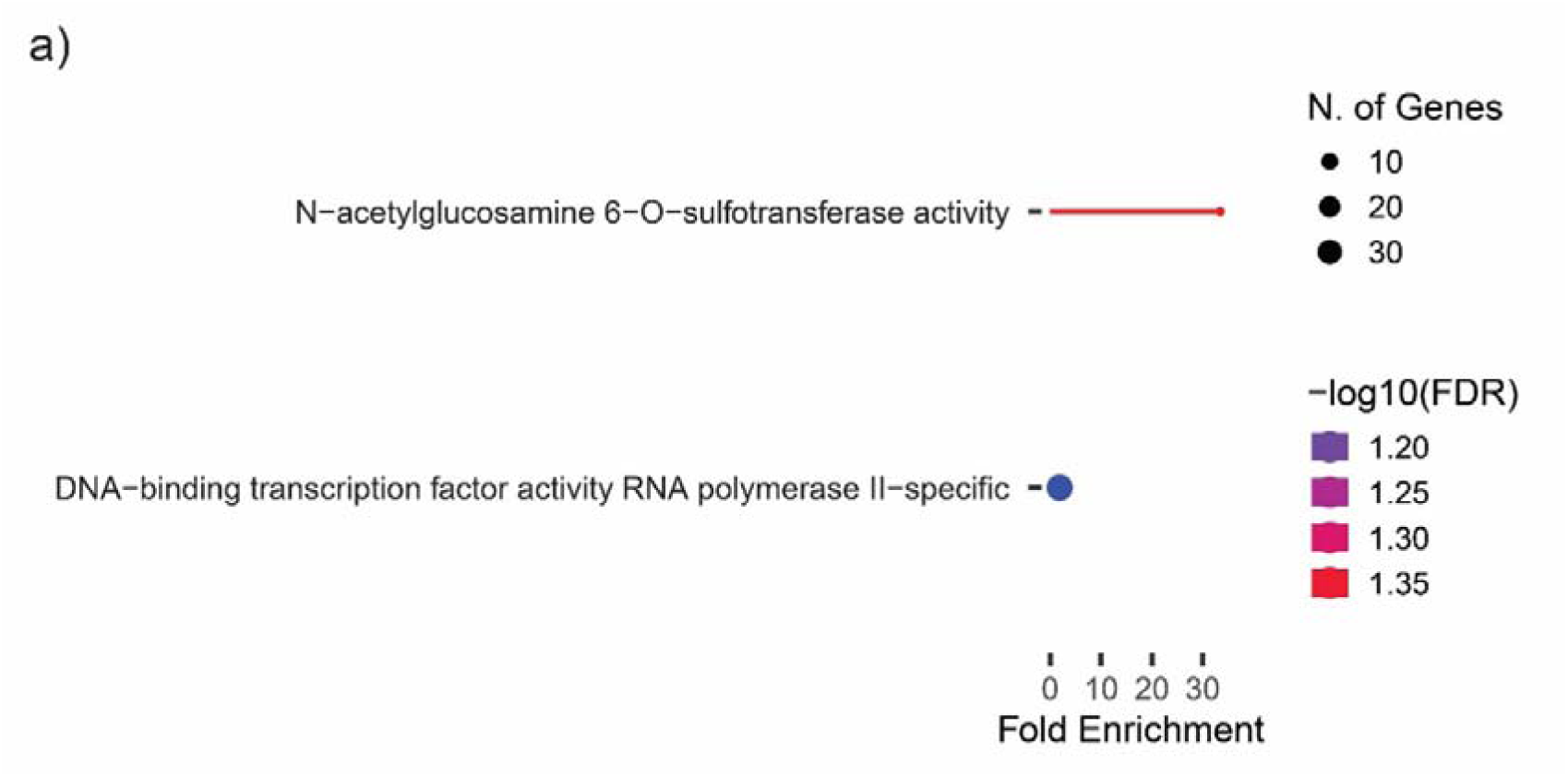
Gene ontology analysis of paraventricular hypothalamic sequencing. (A) Gene ontology enrichment analysis of significant transcripts from sequencing of Djungarian hamster paraventricular hypothalamus across 32-weeks of short photoperiod exposure.

**Fig. S6.**
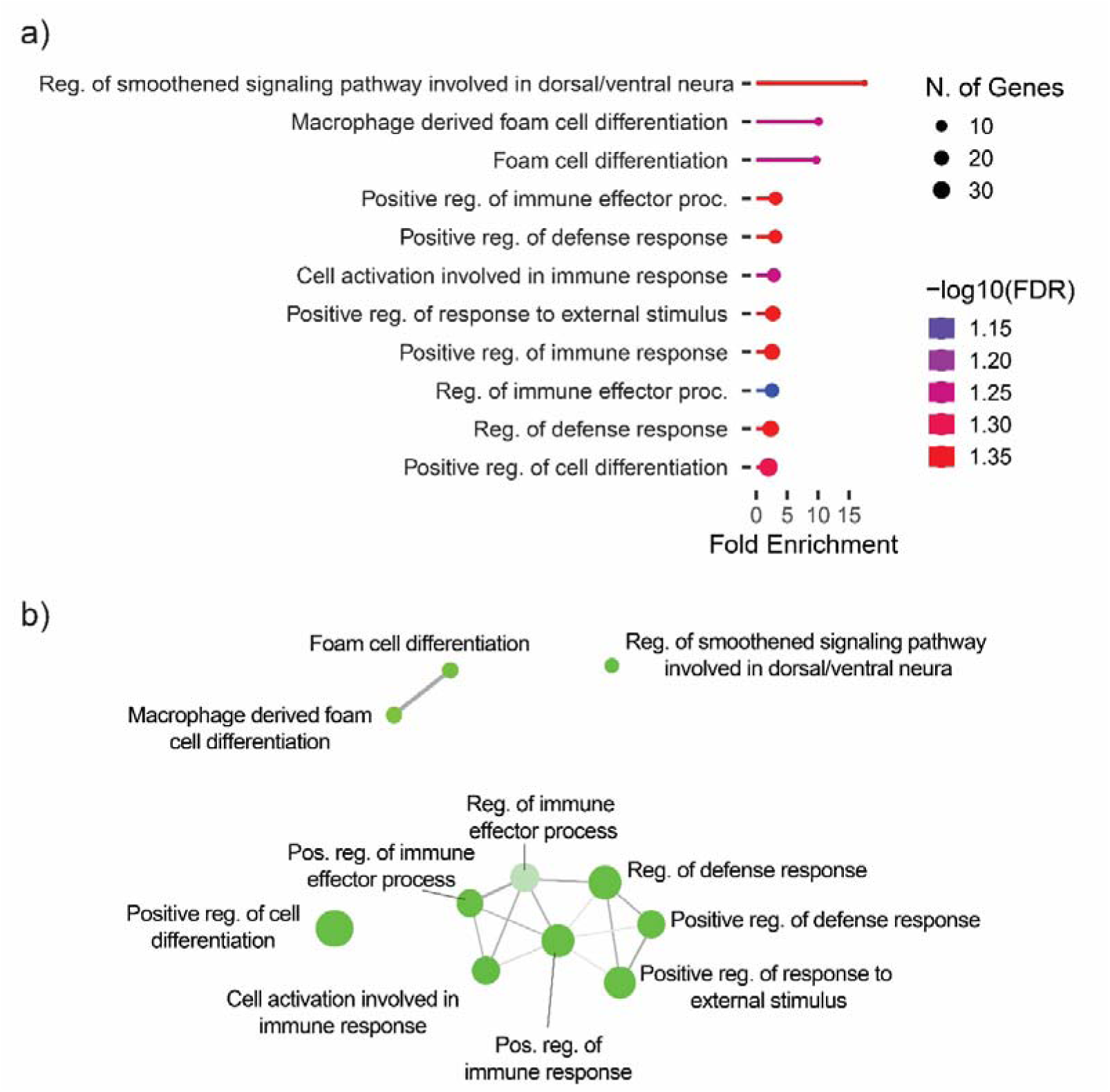
Gene ontology analysis of dorsomedial hypothalamus suggests widespread immune involvement and cellular differentiation. (A) Gene ontology enrichment analysis of significant transcripts from sequencing of Djungarian hamster dorsomedial hypothalamus across 32-weeks of short photoperiod exposure. (B) Interaction network of enriched gene ontology terms. Size of individual points represents the number of transcripts in each term. Thickness of interactions represents shared transcripts between terms.

**Fig. S7.**
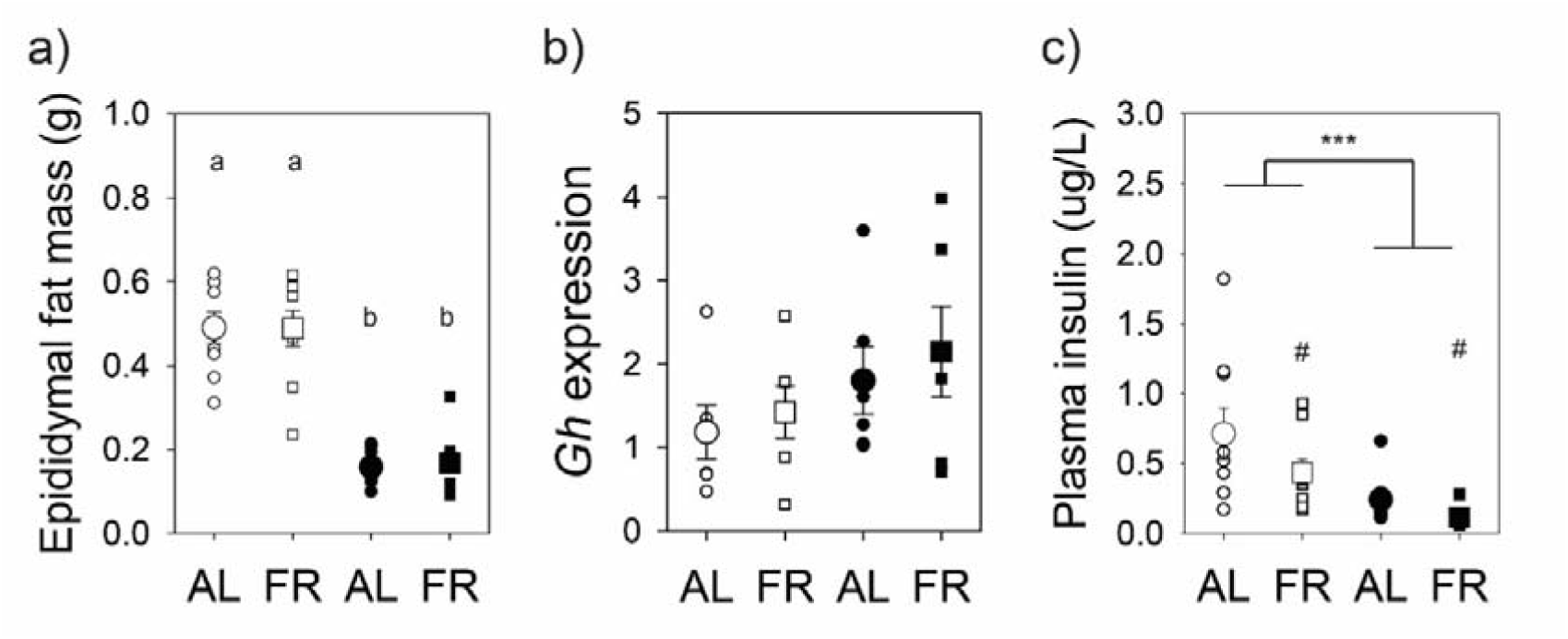
Rheostatic mechanism controlling body mass change in Djungarian hamsters. (A) Epidydimal fat mass in long- and short-photoperiod exposed Djungarian hamsters (F_1,20_ = 59.136; P < 0.001) after an overnight food restriction, or ad libitum, feeding protocol (F_1,22_ = 2.738; P = 0.114). (B) Pituitary gland expression of *Gh* from in long- and short-photoperiod exposed Djungarian hamsters (F_1,20_ = 2.738; P = 0.114) after an overnight food restriction, or ad libitum, feeding protocol, assessed by qPCR (F_1,20_ = 0.518; P = 0.480). (C) Plasma insulin in long-and short-photoperiod exposed Djungarian hamsters (F_1,20_ = 29.865; P < 0.001) after an overnight food restriction, or ad libitum, feeding protocol (F_1,22_ = 6.863; P < 0.05). Letters denote significant main effect of photoperiod (a). Asterisk indicates significant effect of photoperiod (P < 0.005), and hashtag denotes significant main effect of food restriction (P < 0.05).

**Fig. S8.**
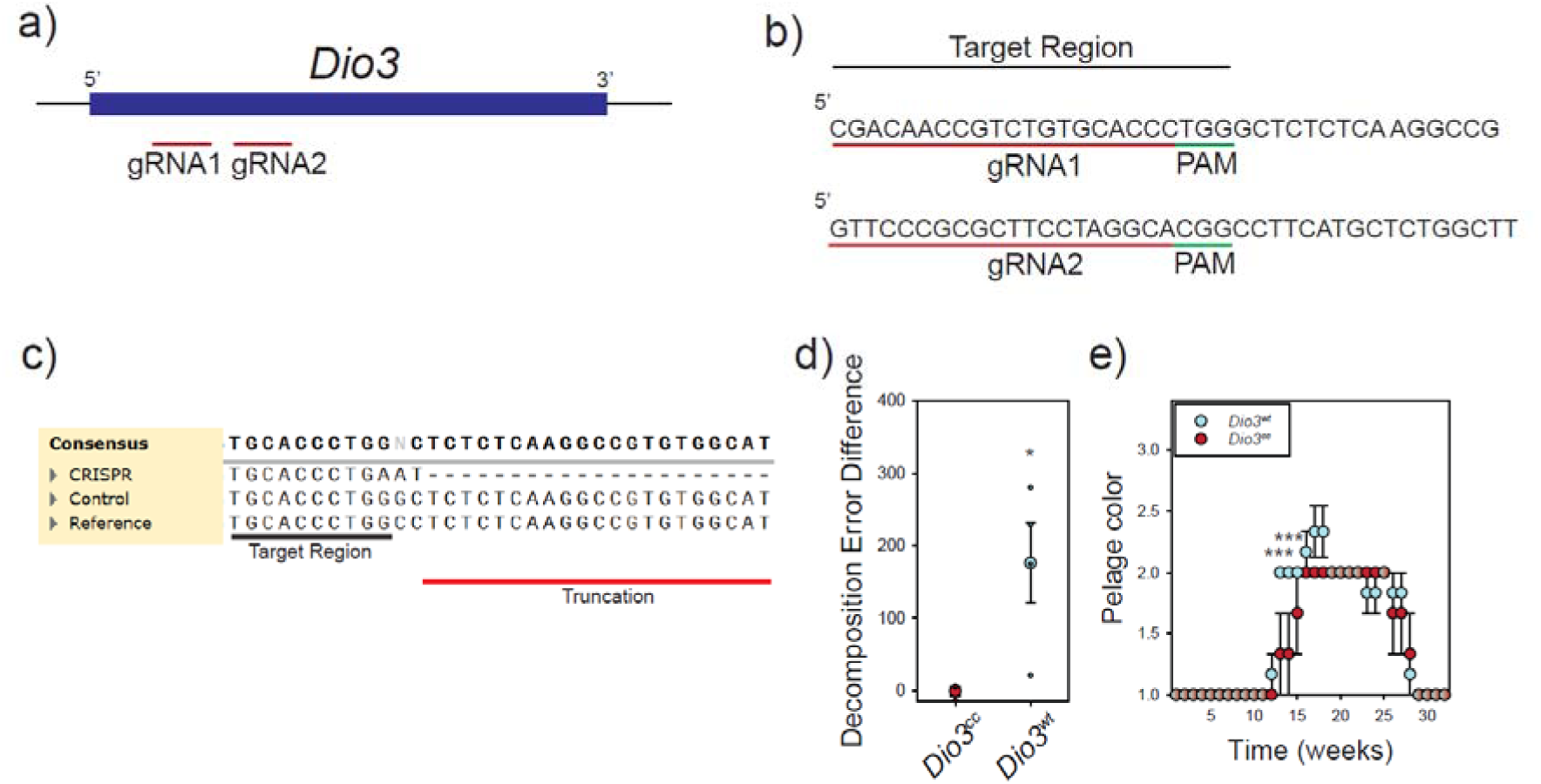
Evidence of CRISPR modification in *Dio3^cc^* and *Dio3^wt^* Djungarian hamsters. (A) Binding positions of gRNA1 and gRNA2, injected into Djungarian hamster hypothalamus (B) Sequence of gRNA1 and gRNA2 binding and PAM sites. (C) Representative *Dio3* sequence alignment with gRNA1 and PAM sites indicated for a representative *Dio3^wt^* (control) and *Dio3^cc^*(CRISPR) hamster. (D) Decomposition error difference for *Dio3* sequence alignment to determine genomic modification (t_5_ = 2.66, p < 0.05) (E) Pelage change in *Dio3^wt^* and *Dio3^cc^* Djungarian hamsters across 32 weeks of SP exposure, *Dio3^wt^*showed significantly increased pelage score at 13 (P < 0.001) and 14 (p < 0.001).

**Table S1.**
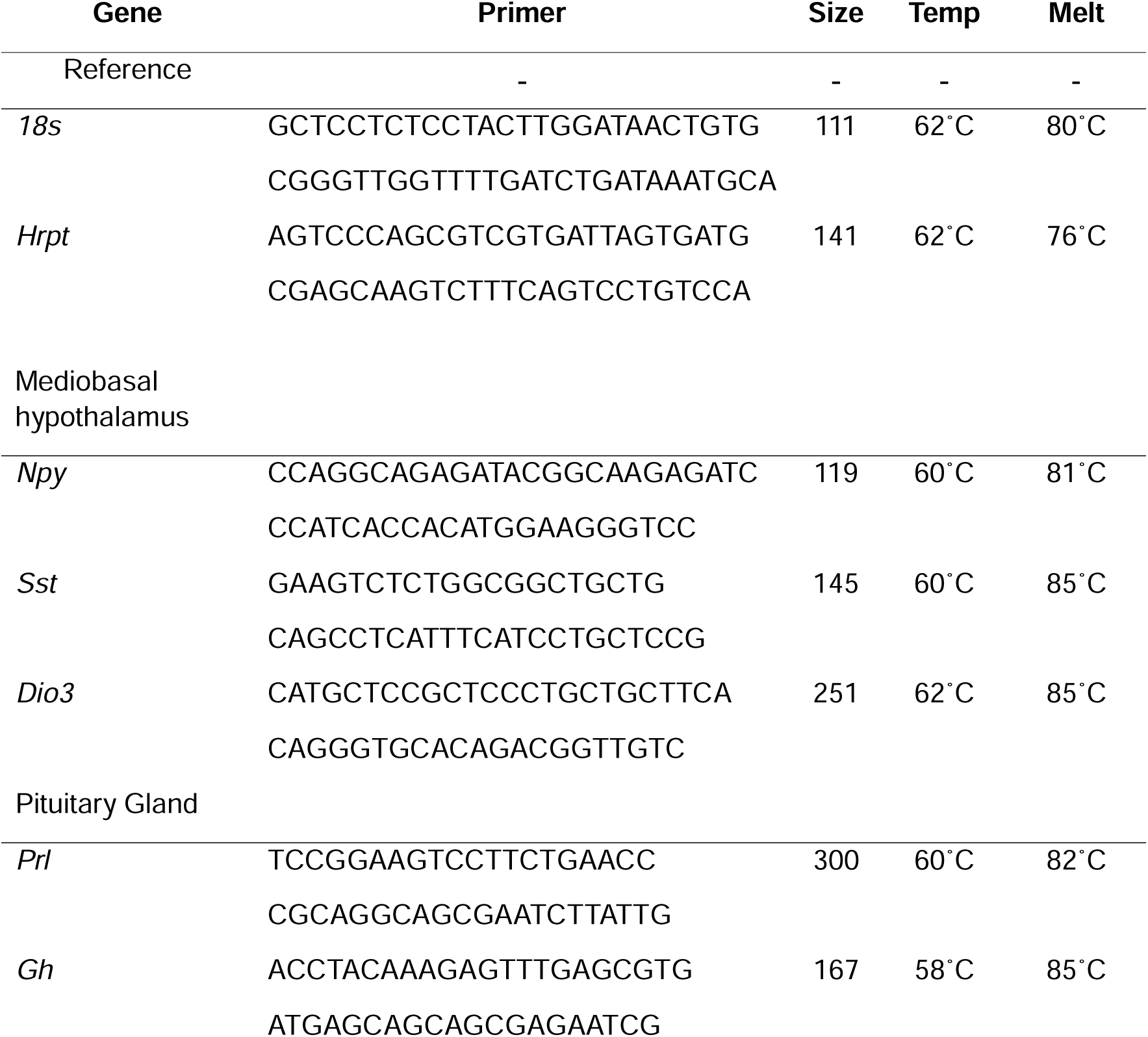
Djungarian hamster qPCR primers sequences and thermal cycling profiles.

## Notes

### Competing Interest Statement

The authors have declared no competing interest.

### Summary of Updates

No major changes to the manuscript. There are minor additions, clarifications or changes in figure number.

