## Supplementary figures and images for "Hypothalamic deiodinase type-3 establishes the period of circannual interval timing in mammals"

### figure S1

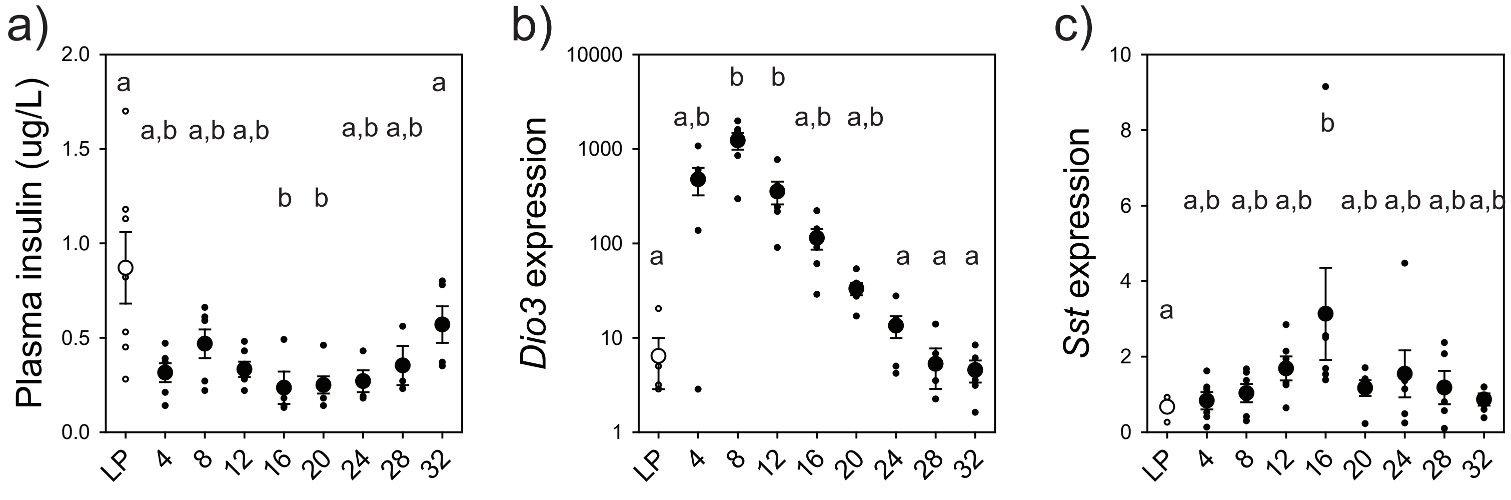

### figure S2

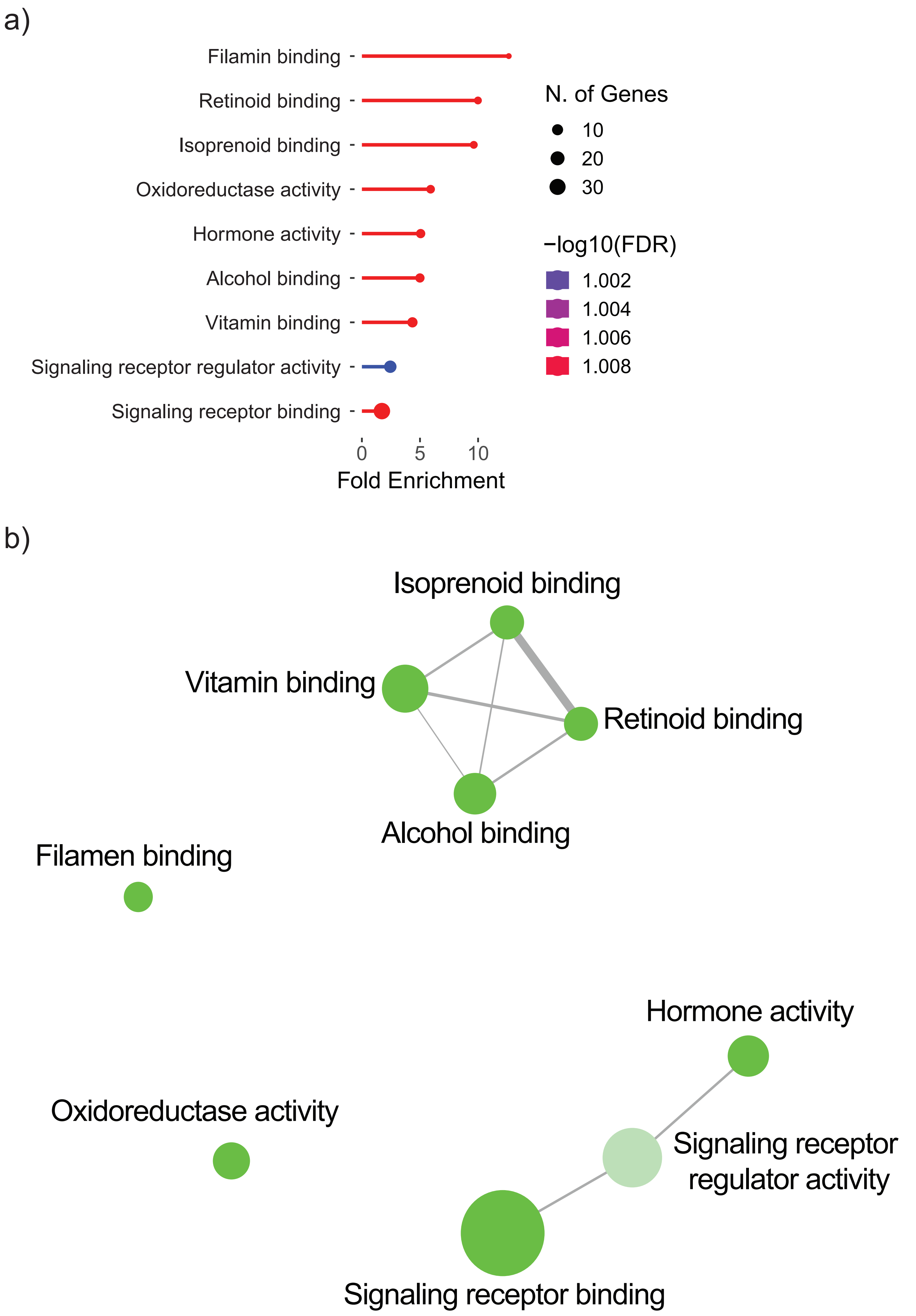

### figure S3

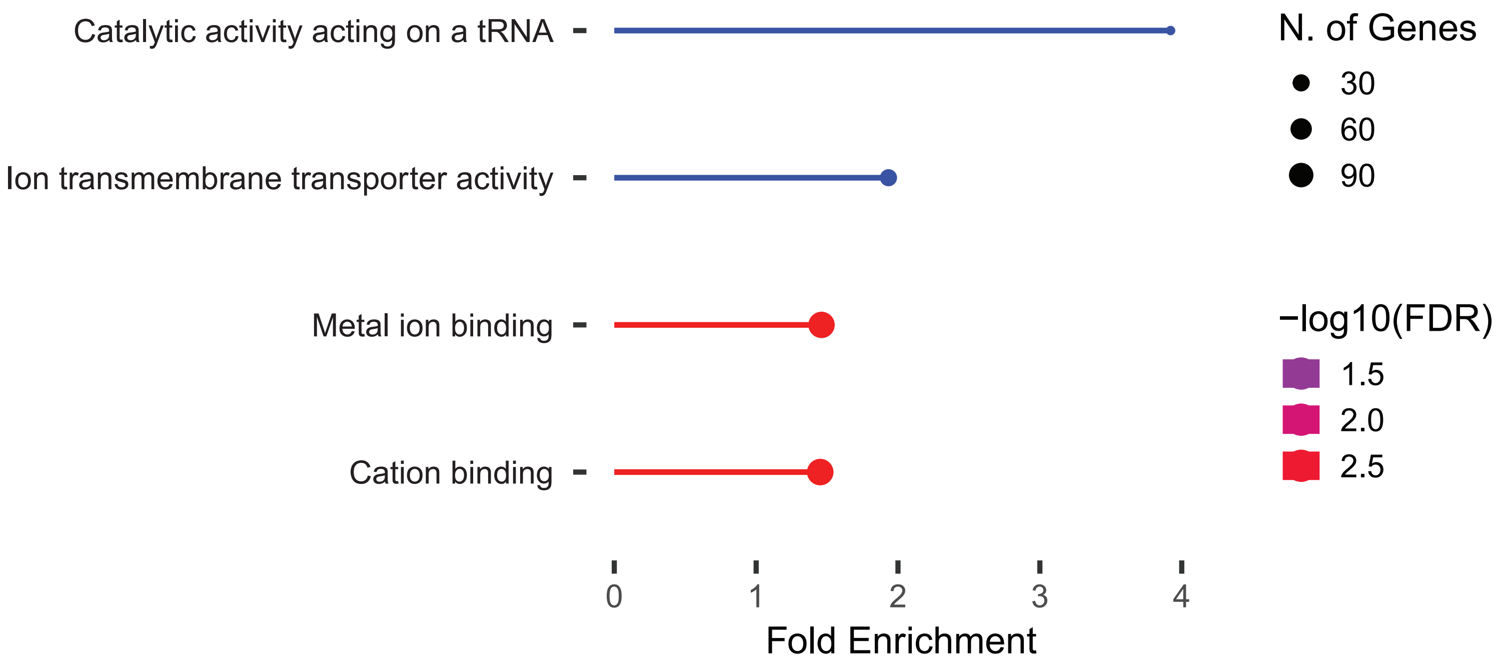

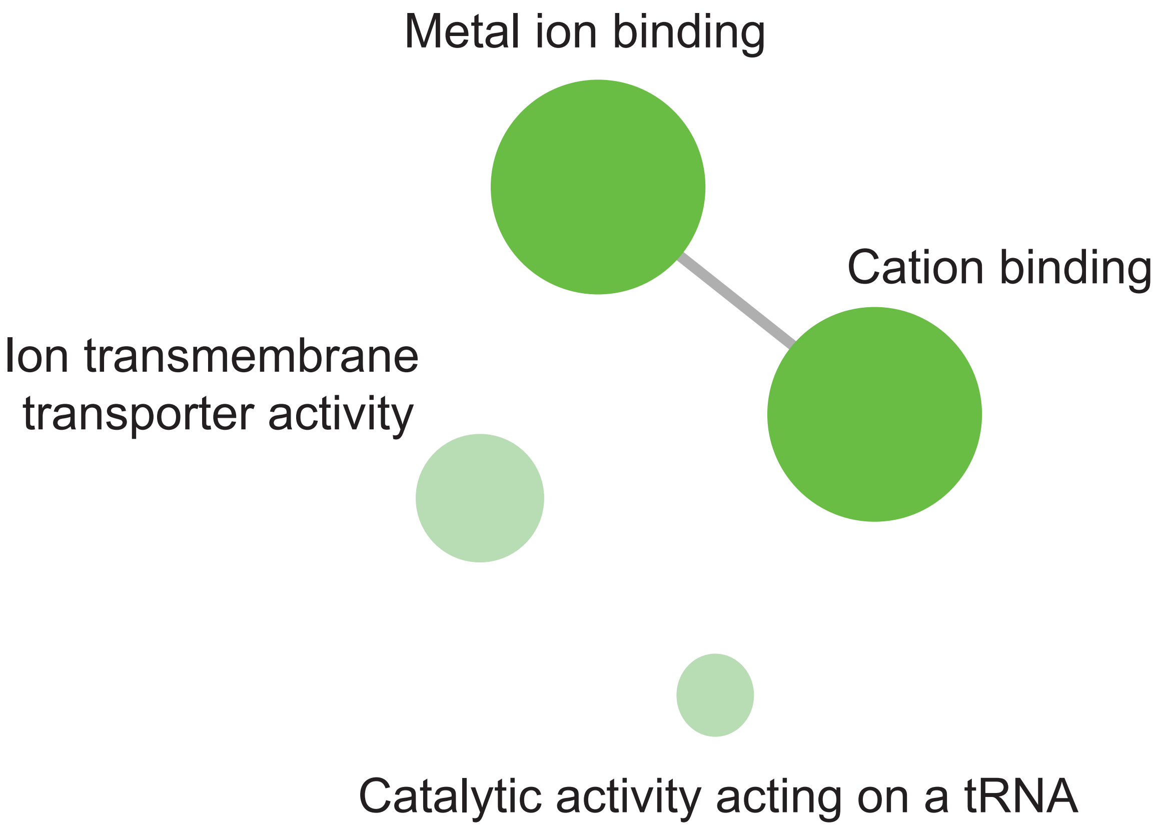

### figure S4

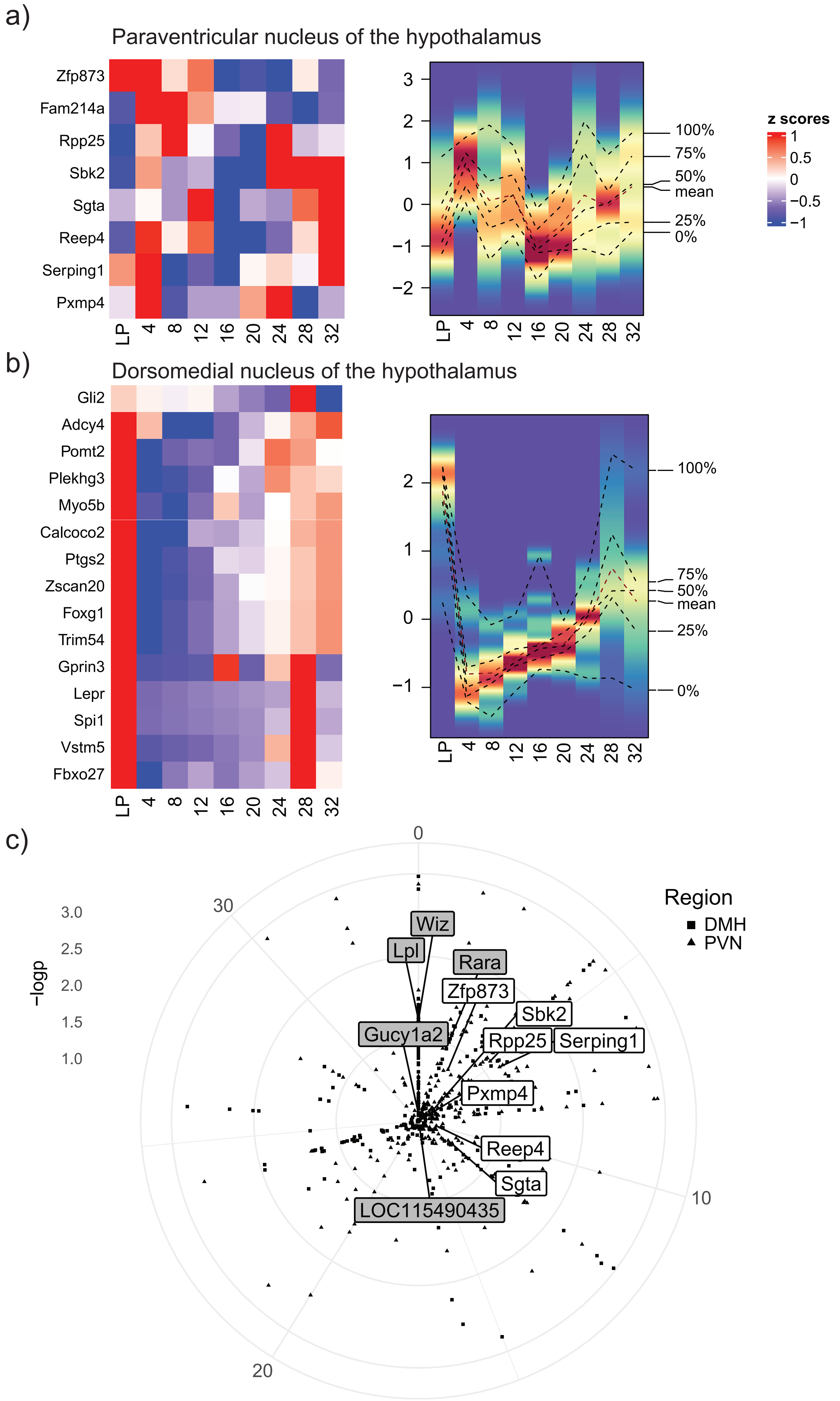

### figure S5

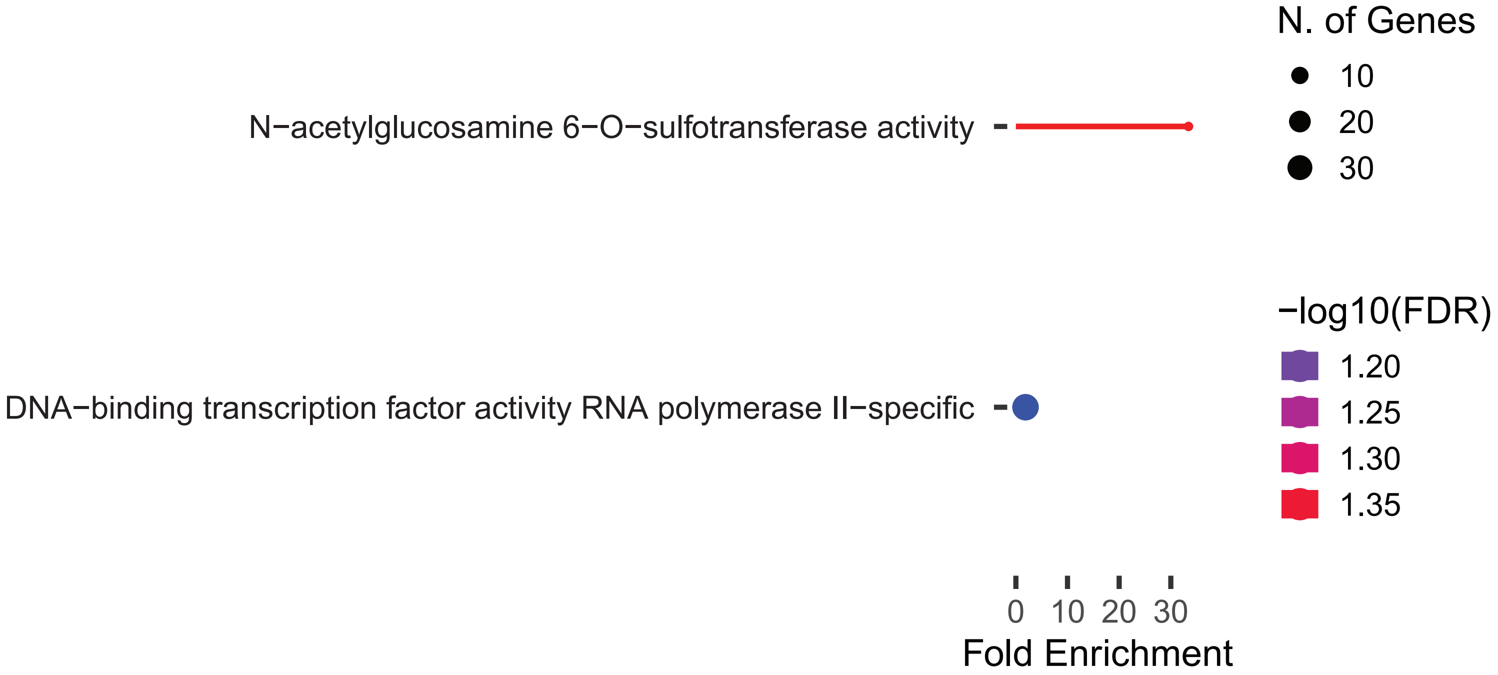

### figure S6

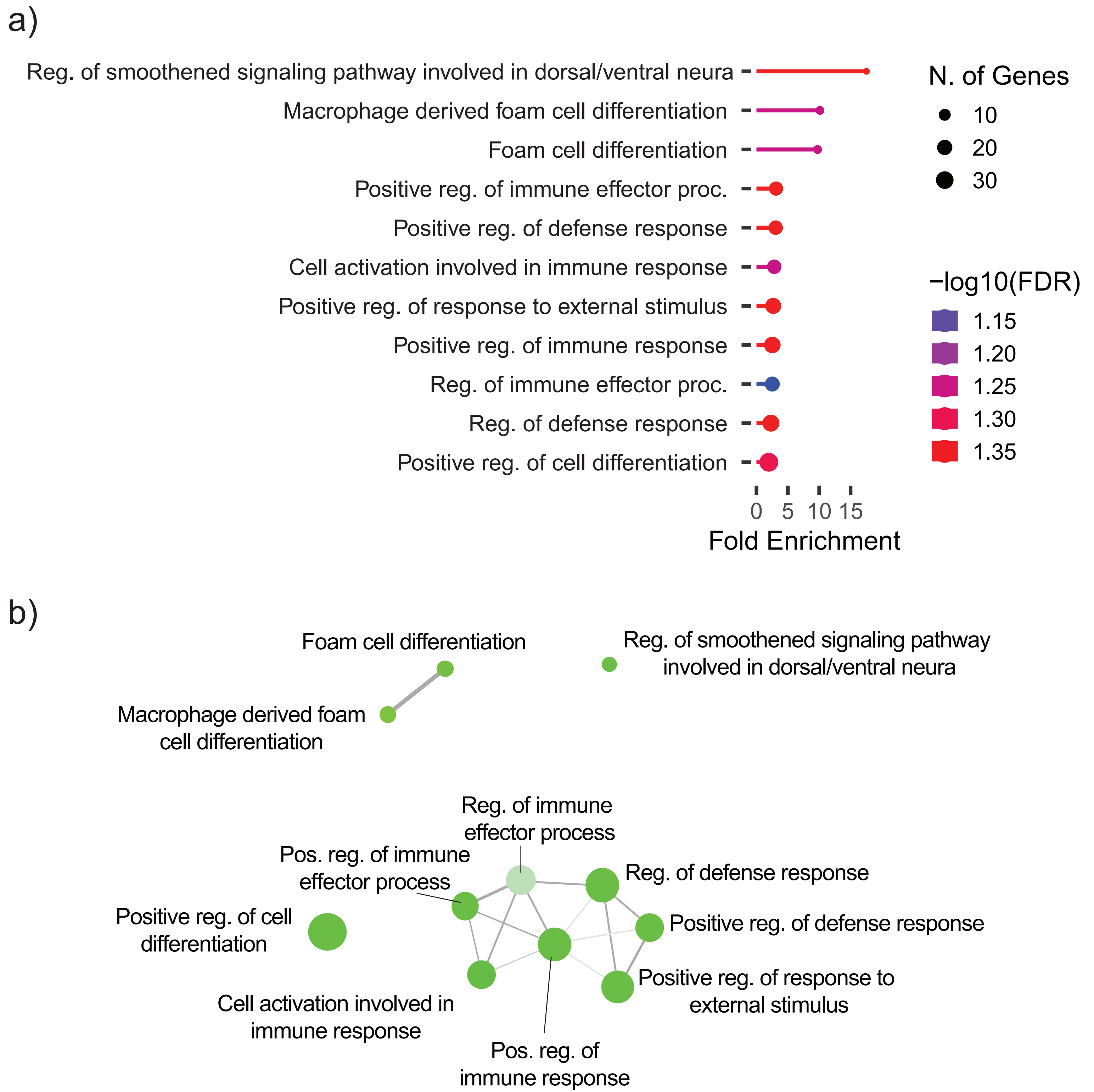

### figure S7

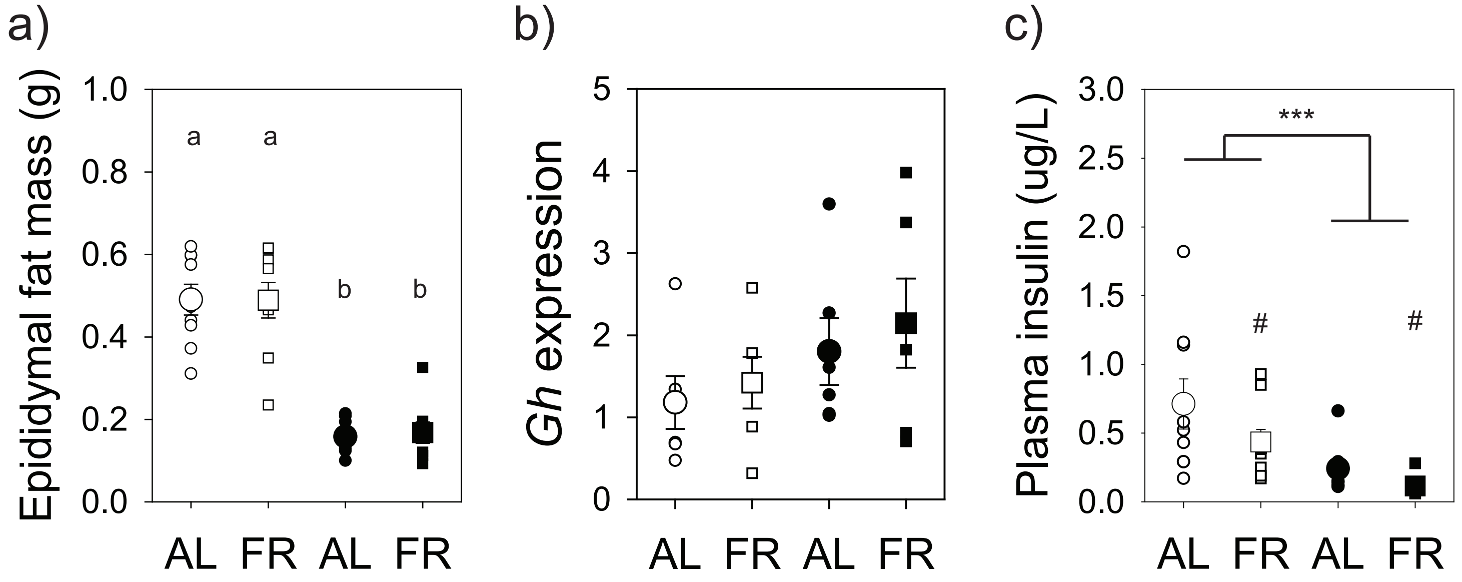

### figure S8

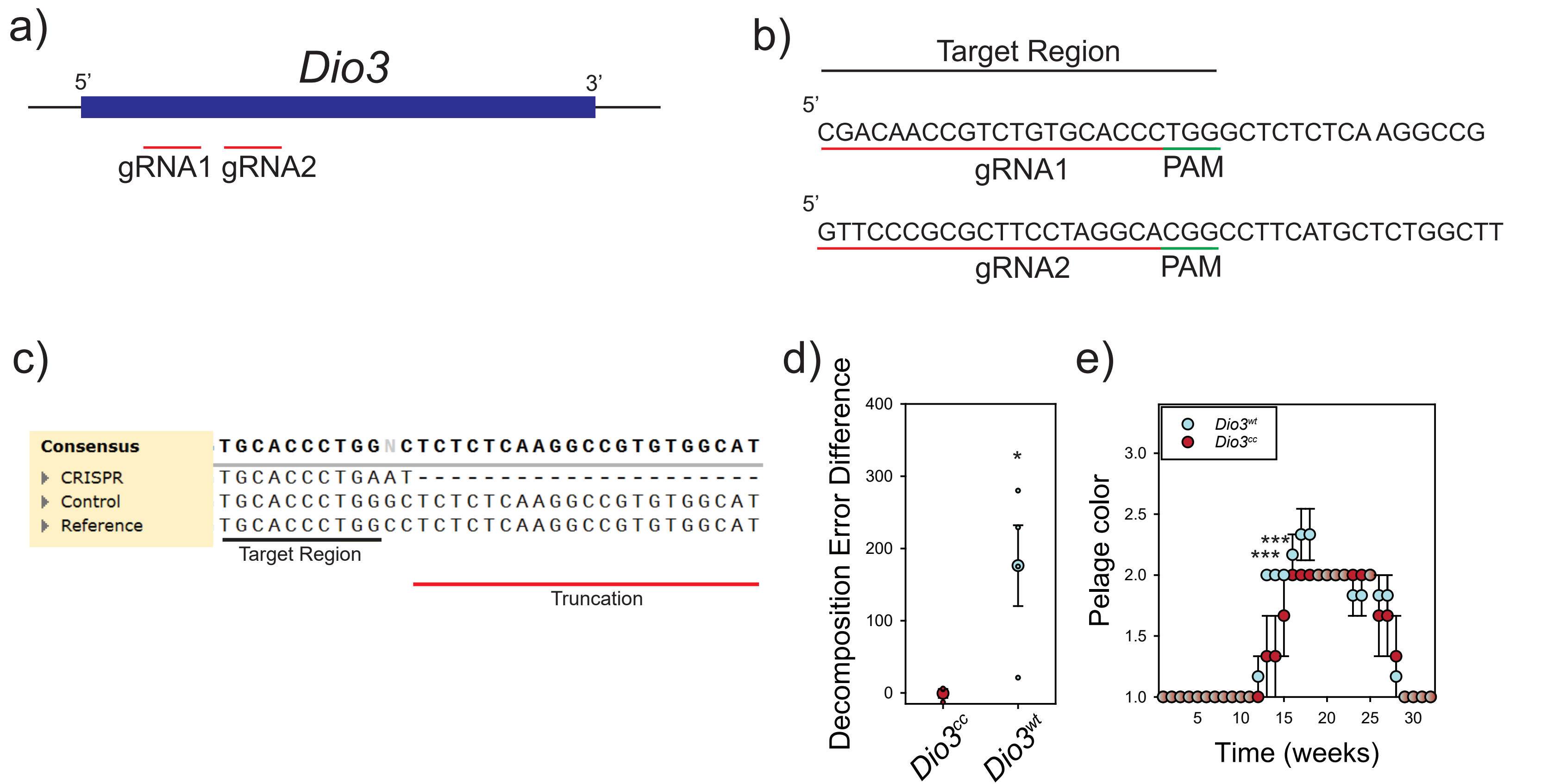
